# Compositional neural dynamics during reaching

**DOI:** 10.1101/2025.09.04.674069

**Authors:** Mehrdad Kashefi, Jonathan A. Micheals, Rhonda Kersten, Jonathan C. Lau, Jörn Diedrichsen, J. Andrew Pruszynski

**Affiliations:** Western Centre for Brain and Mind, Western University, London, Ontario, Canada; Department of Physiology and Pharmacology, Western University, London, Ontario, Canada; School of Kinesiology and Health Science, York University, Toronto, Ontario, Canada; Department of Clinical Neurological Sciences, London Health Sciences Centre, Western University, London, ON, Canada; Department for Computer Science, Western University, London, Ontario, Canada; Department for Statistical and Actuarial Sciences, Western University, London, Ontario, Canada

## Abstract

The complex mechanics of the arm make the neural control of reaching inherently posture dependent. Because previous reaching studies confound reach direction with final posture, it remains unknown how neural population dynamics in the motor cortex account for arm posture. Here we address this gap with high-density neural recordings and a reaching task in which the same targets serve as start points on some trials and end points on others. We show that neural population dynamics in monkey primary motor cortex and dorsal premotor cortex exhibit a compositional structure with three components that enable posture-dependent control: first, a posture subspace containing fixed points visited whenever the arm is in a specific posture; second, rotational dynamics that transition between these fixed points, systematically organized so that similar rotations produce similar movements while continuously updating the posture representation; third, a condition-independent shift dimension that tracks trial progression across all movements. This compositional structure advances the population-level account of how motor cortical dynamics support skilled reaching.

## Introduction

How does the motor cortex generate voluntary reaching movements? A prevalent view is that motor cortical neurons collectively plan and execute reaches through coordinated population activity that unfolds in a low-dimensional neural state space^1,2^. During movement preparation, the neural population state is set to an initial condition that then evolves during execution to drive spinal circuits and produce the appropriate muscle activity patterns^3–7^. This view successfully links movement preparation and execution^8,9^, describes how the motor cortex contributes to decision making^10,11^, anticipates reward^12,13^ and adjusts to sensory feedback^14,15^, and reveals the neural basis of movement corrections^16,17^ as well as sequential actions^18^.

An important issue that remains unsolved is how motor cortical dynamics are organized for movements at different postures. Addressing this problem is critical since the control policy for a complex effector like the arm is inherently posture dependent^19–21^. Consistent with this biomechanical reality, posture-related signals are abundant in motor cortical areas. Many single neurons are tuned to hand position^22–27^ or joint configuration^28–32^, and even direction-tuned neurons are often sensitive to arm configuration^33,34^. At the population level, posture signals occupy a dedicated subspace during preparation that remains orthogonal to dimensions that code movement goals^35,36^. What remains elusive is a unified understanding of how motor cortical population dynamics support posture-dependent control. This question remains unanswered because of the widespread reliance on center-out reaching tasks^3,4,6–9^. By initiating all movements from a single starting location, center-out tasks conflate movement direction with the arm*’*s final posture. This makes it impossible to disentangle neural dimensions related to movement dynamics from those related to postural state, as the two always evolve together during movement.

We disentangled posture and control signals by training monkeys to perform delayed reaches between all possible pairs of five targets. Critically, each target served as a start location on some trials and an end location on others, enabling us to assess the neural state associated with postural state independent of movement dynamics. High-density recordings from primary motor cortex (M1) and dorsal premotor cortex (PMd) revealed neural population dynamics with a compositional structure^37^ featuring three key parts. First, a posture subspace contained fixed points for each spatial target and was visited whenever the arm rested at that target before or after a reach. Second, a set of rotational dynamics linked these fixed points during movement. Movement dynamics were organized such that more similar rotations were associated with more similar reach directions, and their projection onto the posture subspace provided continuous information about arm posture. Third, a condition-independent shift dimension reflected trial progression, which unfolded similarly across conditions regardless of movement parameters. These findings provide a more complete account of motor cortical geometry, showing how neural population activity simultaneously generates movement, maintains an internal representation of limb posture, and tracks task progress.

## Results

### Task, behaviour, and recording setup

We trained two rhesus macaques (Monkey M and Monkey P) to perform delayed reaching movements using a KINARM^38^ exoskeleton robot (**Fig. 1a**). The monkeys reached between all possible pairs of five targets positioned at the corners and center of a 10 × 6 cm rectangle (**Fig. 1b**). Crucially, each target served as a start point on some trials and an end point on other trials which allowed us to observe the neural state associated with postural state independent of movement dynamics. This design also allowed us to disentangle the contributions of movement parameters. For example, some diagonal reaches shared the same start location and direction but differed in extent (e.g., cross to square vs. cross to diamond). Other reaches shared similar direction and extent but differed in start and end locations (e.g., cross to plus vs. circle to diamond) (**Fig. 1b**). Each trial began with the presentation of one of the five targets as the start location. After the monkey held its hand at the start position for 500 ms, one of the remaining four targets was randomly selected and displayed as the goal target. The monkey was required to remain stationary at the start target for a variable delay period (300–700 ms) until a go cue—an auditory beep—signaled the start of movement (**Fig. 1c**). Monkeys initiated the reach from the start target promptly after the go cue and quickly and accurately moved their hand to the end target (**Fig. 1d**). Both monkeys maintained a high success rate (>90%) across sessions. We used 45mm NHP Neuropixels probes to record neural activity from the primary motor cortex (M1) and dorsal premotor cortex (PMd) in both monkeys. In Monkey P, we additionally recorded from primary somatosensory cortex (S1), dorsolateral prefrontal cortex (dlPFC, Brodmann area 46), supplementary motor area (SMA), pre-supplementary motor area (pre-SMA), and the internal segment of the globus pallidus (GPi) (**Fig. 1e; see Methods**).

**Figure 1.**
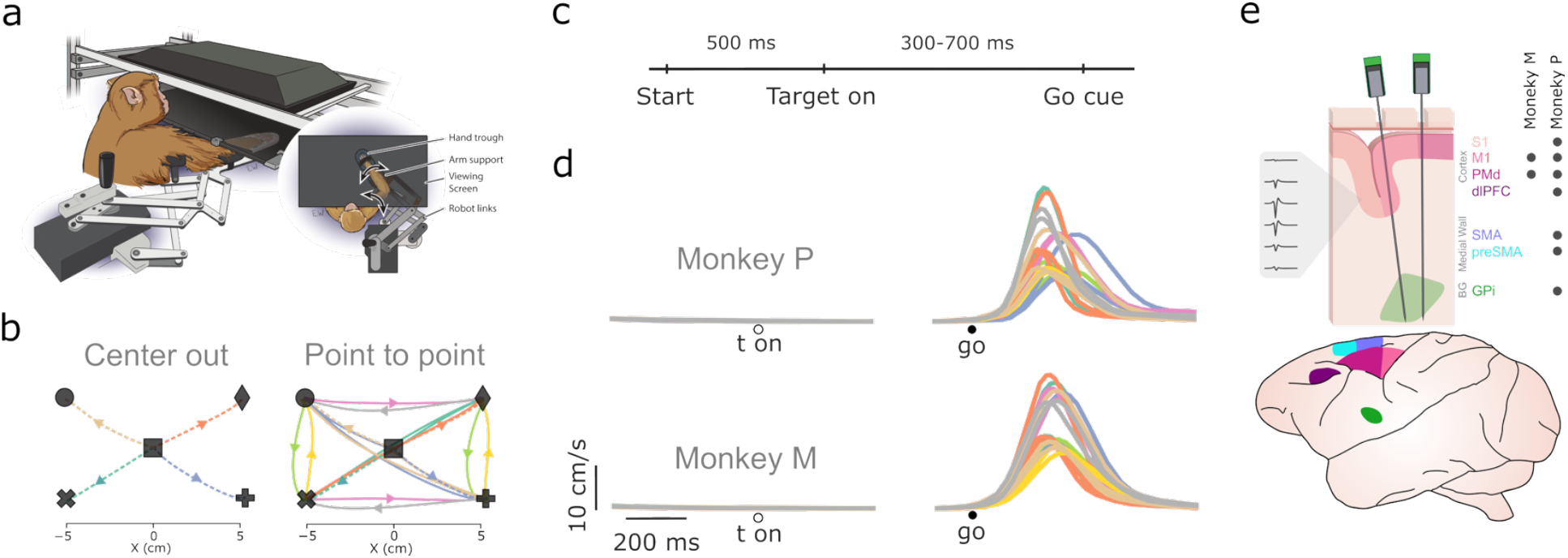
Setup and behaviour. **a)** KINARM exoskeleton setup. **b)** Potential reach targets on the center and vertices of a rectangle. NHPs performed center-out reaches as well as reaches between all possible pairs of targets – all conditions were randomly interleaved. Each trace is the average hand path for monkey P, colored by reach direction. Note that although we plot targets at different positions using different symbols for semantic clarity, all targets presented to the monkey were identical. **c)** Timing of the task. Each trial had a new start target; the end target was shown when NHPs remained in the start target for 500 ms. **d)** Average speed profile for all twenty conditions colored by reach direction for Monkey P and Monkey M. **e)** Schematic of Neuropixels 1.0 NHP Long and areas recorded in Monkey P and Monkey M.

### Majority of single units show mixed selectivity to movement parameters

Consistent with previous studies, single neurons in motor cortex generally showed mixed selectivity^39,40^ and complex temporal activity patterns^33,41,42^. Although we found some neurons that were predominantly active during the preparation phase (**Fig. 2a, top**) and others that were primarily active during movement execution (**Fig. 2a, bottom**), the majority exhibited complex and heterogeneous activity patterns across both epochs^43,44^. To quantify the strength and frequency of parameter selectivity in our task, we analyzed each recorded neuron separately during the preparation and execution epochs. For each epoch, a neuron was considered selective for start position (P), direction (D), or extent (E) if a classifier could predict that parameter from the neuron*’*s firing rate significantly above chance (**see Methods**). Based on decoding performance, neurons were categorized as selective for a single parameter (P, D, or E), a combination of two parameters (PD, DE, or EP), or all three simultaneously (PDE) (**Fig. 2b**).

**Figure 2.**
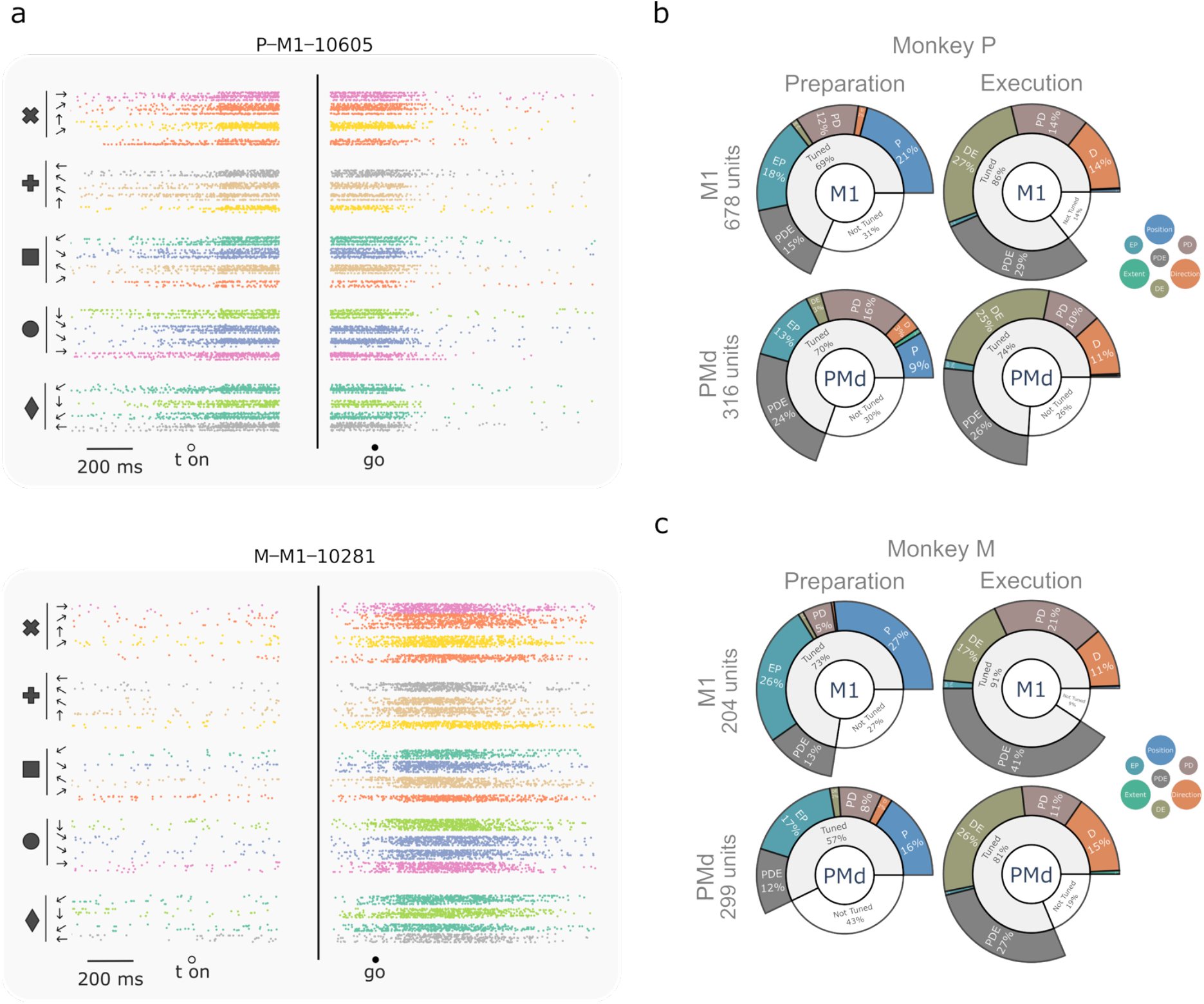
Majority of single neurons show mixed selectivity. **a)** Two example M1 neurons. Raster is aligned on target onset (left), and on go cue (right). Trials are grouped by start position (shapes) and movement direction (arrows, colors). **b)** Fraction of tuned units for M1 and PMd recordings from Monkey P during preparation (left), and execution (right) phase of the movement. For the tuned single neurons, each neuron is further categorized based on their tuning to start Position (P, blue), movement Direction (D, orange), and movement Extent (E, green), as well as combinations of these parameters (shown in mixed hues). **c)** Same as (b) for Monkey M.

In Monkey P*’*s M1, during the preparation phase, 21% of all isolated neurons were tuned to start position, 2% to direction, and the majority exhibited mixed selectivity for multiple movement parameters. During execution, only 14% of neurons were tuned to direction, while the remaining neurons continued to show mixed selectivity (**Fig. 2b**). A similar pattern was observed in Monkey P*’*s PMd, as well as in Monkey M*’*s recordings from both M1 and PMd (**Fig. 2c**). Tuning distributions from additional areas recorded in Monkey P are shown in Supplementary Fig. 1. Consistent with widespread mixed selectivity at the single-neuron level, the population activity collectively encoded all three movement parameters—position, direction, and extent. To determine when these parameters became decodable, we applied a time-resolved classification analysis across the trial, using the full neural population to predict position, direction, and extent at each time point (**see Supplementary Fig. 2 and Methods**). Start position was decodable throughout the entire trial, while direction and extent became reliably decodable during the preparation phase, shortly after target onset.

### Rotational dynamics connect posture-specific fixed points

How do motor cortical areas generate the neural activity needed to transition between postures while simultaneously maintaining an updated representation of limb position? Given the immense heterogeneity and mixed selectivity of single neurons, this question can only be meaningfully addressed at the neural population level. Therefore, we applied principal component analysis (PCA) to M1 population activity. We aligned the neural data and its associated kinematics (**Fig. 3a, left;** position and speed traces) to the *“*go cue*”*, utilizing a window spanning –700 ms to +1400 ms relative to the cue. This window captured the resting period at the start target (before target onset), the preparation phase, and the full execution of the reach (**Fig. 3a, left;** see Methods).

**Figure 3.**
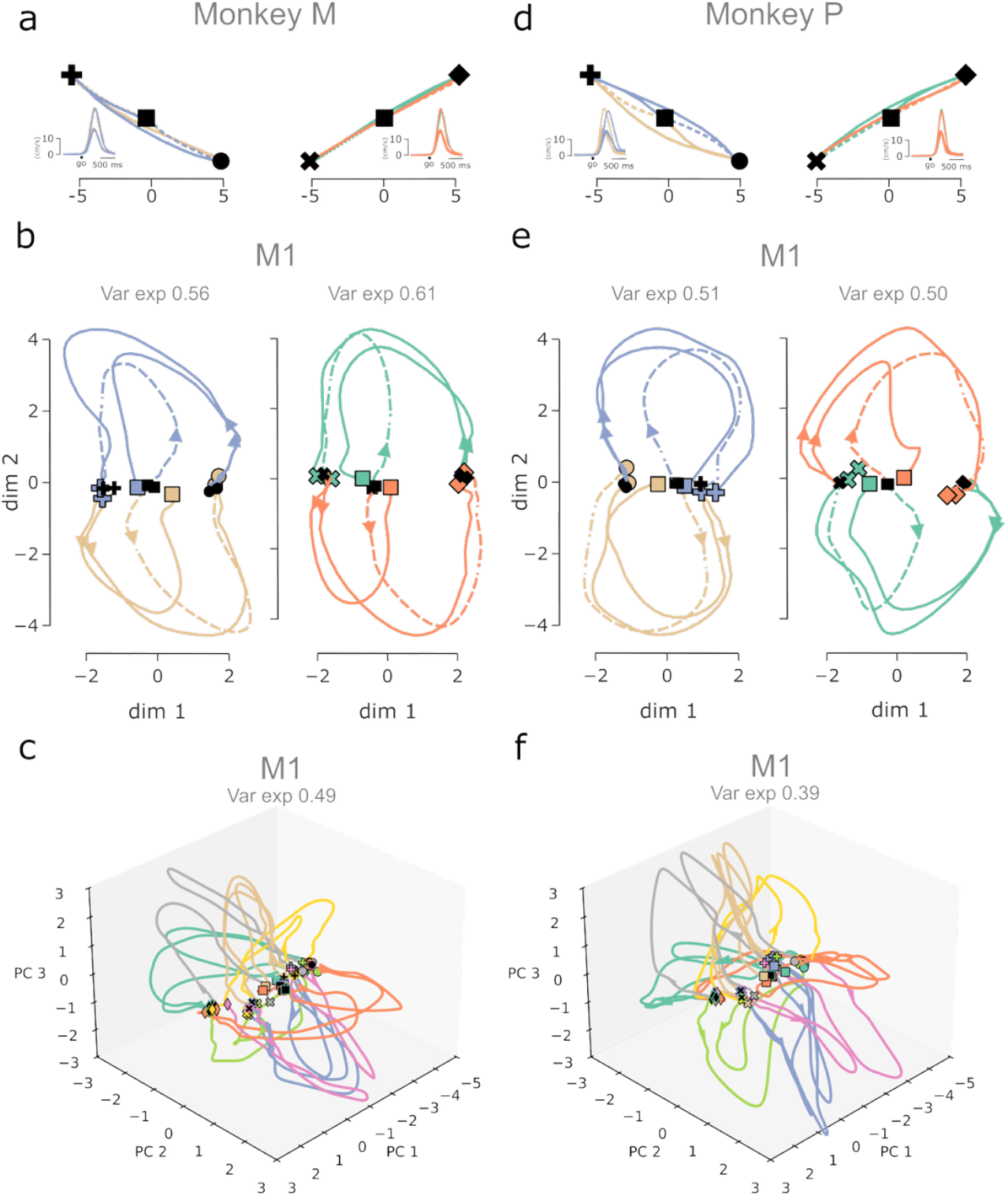
Rotational dynamics connect posture-specific fixed points during reaching. **a)** Hand trajectories and speed profiles for six trial types involving reaches between three diagonal targets. Colors indicate reach direction. **b)***Left:* M1 Neural trajectories for each reach condition. Dotted traces represent center-out reaches. Each trace begins at a black marker indicating the start location, shaped as a plus, square, or circle corresponding to trials that began from the plus, square, or circle target, respectively. Traces are colored by reach direction, and arrows indicate the go cue. Each trace ends in a colored marker (plus, square, or circle) representing the end target location. *Right:* Same as left, but for the trial types shown in panel(a) (right). **c)**Neural trajectories projected onto the top three principal components (PCs) from a PCA fit across all 20 conditions. Each trace represents one reach condition, colored by reach direction. As in panel (b), black markers indicate start locations and colored markers indicate end locations. See this figure online. **d–f)** Same as panels a–c, but for Monkey P. See this figure online.

We first examined two canonical center-out conditions: *square* → *plus* and *square* → *circle* (**Fig. 3a**, left, dashed traces). In both cases, the neural state began at the same location in state space (**Fig. 3b, left;** black square). During the preparation phase, the states diverged along dim 2, and at the go cue (arrow), they rotated in opposite directions toward distinct endpoints on dim 1(**Fig. 3b**). Crucially, our design allowed us to ask whether the neural state for a reach *starting* from a given posture matched the neural state where a reach to that same posture *ended*—that is, whether the same neural states were reused for out-center and center-out movements. Indeed, we found that neural states were reused. In a *plus* → *square* reach, the neural state began where the *square* → *plus* reach ended, then during preparation detached from dim 1 and, at the go cue, evolved along a rotational trajectory toward a location close to the initial state of reaches starting from the square posture. Looking at all conditions including these three targets, the same pattern emerged with the neural state transitioning between three fixed points as the arm moved between all three target locations. We observed the same organization when PCA was fit to reaches along the other diagonal (**Fig. 3a right, Fig. 3b right**) or all experimental conditions reaching between five start locations. Visualizing the geometry for five start locations and their fixed points required at least three dimensions (**Fig. 3c**).

As an alternative view, and to further deconstruct these overlapping signals, we identified three mutually orthogonal subplaces optimized to capture variance associated specifically with posture, preparation, or execution (**Supplementary Fig. 3;** see Methods). The dynamics unfolding in these subplaces were distinct: Activity in execution subspaces was more rotational compared to posture and preparation subspaces (**Supplementary Fix. 4;** see Methods). Crucially, as expected from the simple PCA analysis, the isolated posture subspaces continuously represented posture from the start to the end of the trial. This continuous update occurred simultaneous with transitional activity related to preparation and execution occurring in their respective subspaces (**Supplementary Fig. 3**).

Overall, both low dimensional views of the data indicate that at rest, the geometry of neural dynamics in the primary motor cortex is primarily organized by posture. Preparation and execution activity occur as momentary transitions between postural fixed points, structured so that their projections onto the posture subspace continually update and track the current posture. These transition dynamics are organized within the neural state space such that dynamics generating movements in similar direction are positioned close to one another (**Supplementary Fig. 5**). We observed the same structure in the second monkey*’*s primary motor area (**Fig. 3d-f**) and in dorsal premotor area of both monkeys. In monkey P, similar dynamics were also observed in SMA, GPi, and S1, but were absent in the pre-SMA and dlPFC (**Supplementary Fig. 6**).

### Geometry of neural state space at start posture

The PCA analysis revealed five distinct fixed points, each associated with one of the five possible start locations. Next, we characterized the geometric arrangement of these fixed points in neural state space. Specifically, we considered two hypothetical geometries. If the neural representation reflects the spatial layout of the targets in the external workspace, then start locations that are physically closer together should have more similar neural representations. Such a geometry could either mirror the Cartesian distances between targets, or the differences in joint coordinates. Because the joint model led to very similar prediction within our relatively small workspace (**Supplementary Fig. 7**), we concentrated here on the Cartesian arrangement (**Fig. 4a**, Cartesian). Alternatively, targets could be represented as unique locations, independent of their physical target location. In this scheme, each target is represented by unique neural state, equidistant from all other target state — akin to an abstract identity code along orthogonal dimension (**Fig. 4a**, Identity).

**Figure 4.**
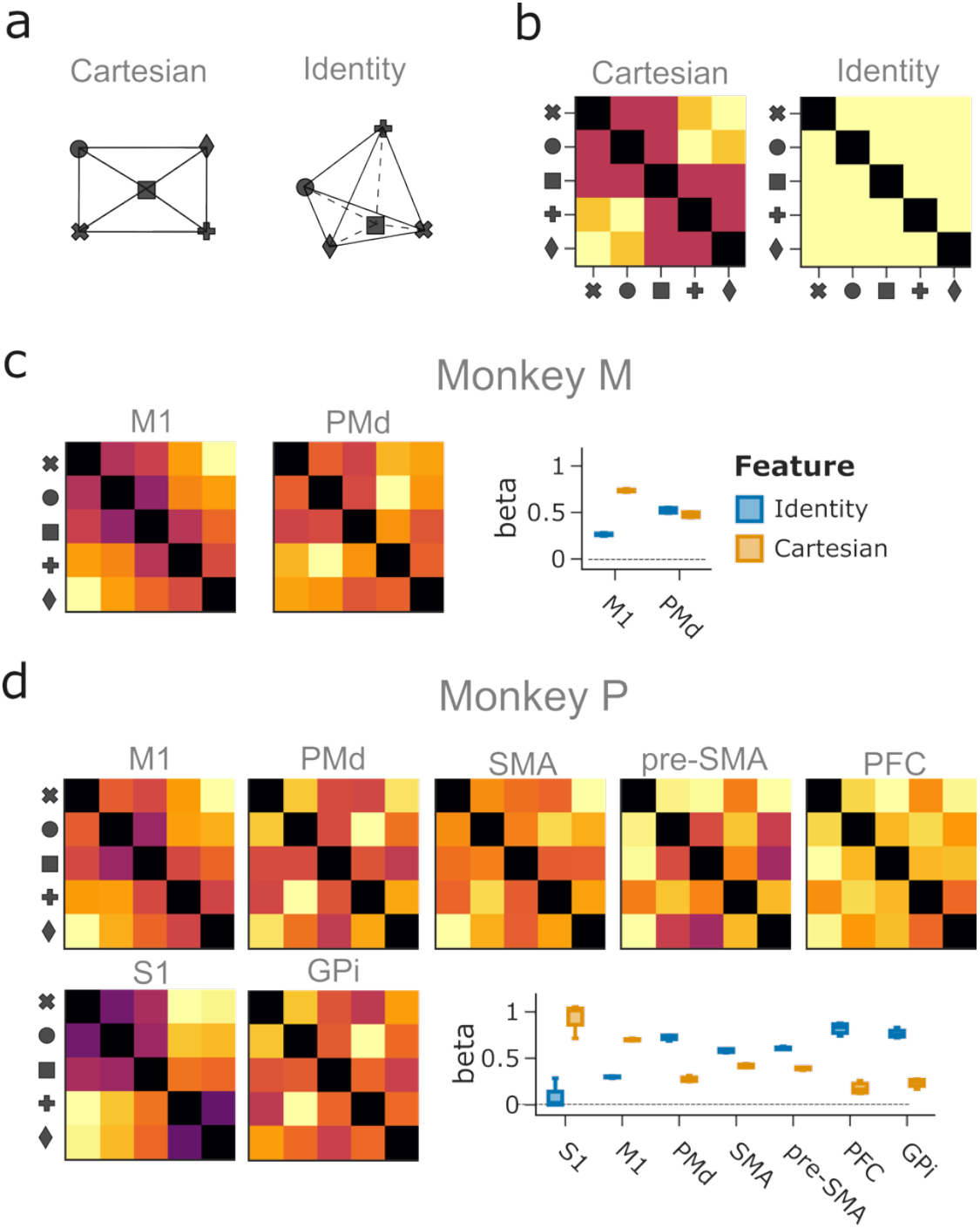
Geometry of neural state space at the start posture. **a)** Two hypothetical geometrical arrangements of fixed points in neural state space. Black lines indicate distances between fixed points. In the *Cartesian geometry*, distances between neural fixed points mirror the spatial distances between targets in the physical workspace. In the *Identity geometry*, all five fixed points are equidistant from one another, representing a fully abstract coding scheme. Note that at least four dimensions are required to position five equidistant points, so the figure shows a 2D projection of this higher-dimensional geometry. **b)** Representational dissimilarity matrices (RDMs) for the Cartesian and Identity models. RDMs for neural data corresponding regression beta weights for the Cartesian and Identity model **c)** RDMs in M1 and PMd of Monkey M. Beta weights indicate each model*’*s contribution to the total variance explained in the regression. **d)** Same as (**c**) but for all recorded regions in Monkey P.

To evaluate the extent to which each hypothesized geometry explains the neural data, we first constructed representational dissimilarity matrices (RDMs) for both the Cartesian and Identity models (**Fig. 4b**). Next, we built an RDM from the neural data by computing the pairwise Euclidean distances between the average firing rates (across all recorded neurons) corresponding to each start location. This was done using the 100 ms window prior to target onset, during which the monkey*’*s hand was stationary at each target. We then used non-negative linear regression to quantify how well the neural geometry could be explained as a combination of the Cartesian and Identity geometries (**Fig. 4c–d**; see Methods for details).

As expected, for both monkeys we found a strong contribution of the Cartesian component in M1 and PMd, with M1 exhibiting the most pronounced Cartesian geometry (**Fig. 4c–d; Supplementary Fig. 7** for same results in joint coordinates). Surprisingly, we also observed a substantial contribution of the identity code in M1, indicating that even this low-level motor area can maintain an abstract representation of target identity. In monkey P, regions including SMA, pre-SMA, PFC, and GPi exhibited mixed coding, with contributions from both Cartesian and identity. Notably, all these areas—like PMd—showed a stronger identity code than Cartesian code, with PFC displaying the most prominent identity neural code. At the other end of the spectrum, S1 showed almost no contribution from the identity component, suggesting nearly pure spatial encoding. Overall, the prominence of Cartesian coding decreased progressively in more frontal areas relative to the central sulcus, suggesting a gradient from spatial to more abstract neural code across the sensorimotor hierarchy.

### Geometry of neural state space at the start and end posture

The PCA projections (**Fig. 3**) indicate that fixed points associated with each posture are reliably engaged whenever the hand rests at a given target—whether at the start or end of a trial. We next asked whether this structure persists in the full dimensional neural state space. If the same fixed point is used at both time points, two predictions follow. First, the distances between fixed points should be similar whether computed from the beginning or end of the trial. Second, for each of the five targets, the neural state at the start of the trial should closely match the neural state at the same target at the end of the trial, yielding near-zero pairwise distances (**Fig. 5a– b**, *Aligned*).

**Figure 5.**
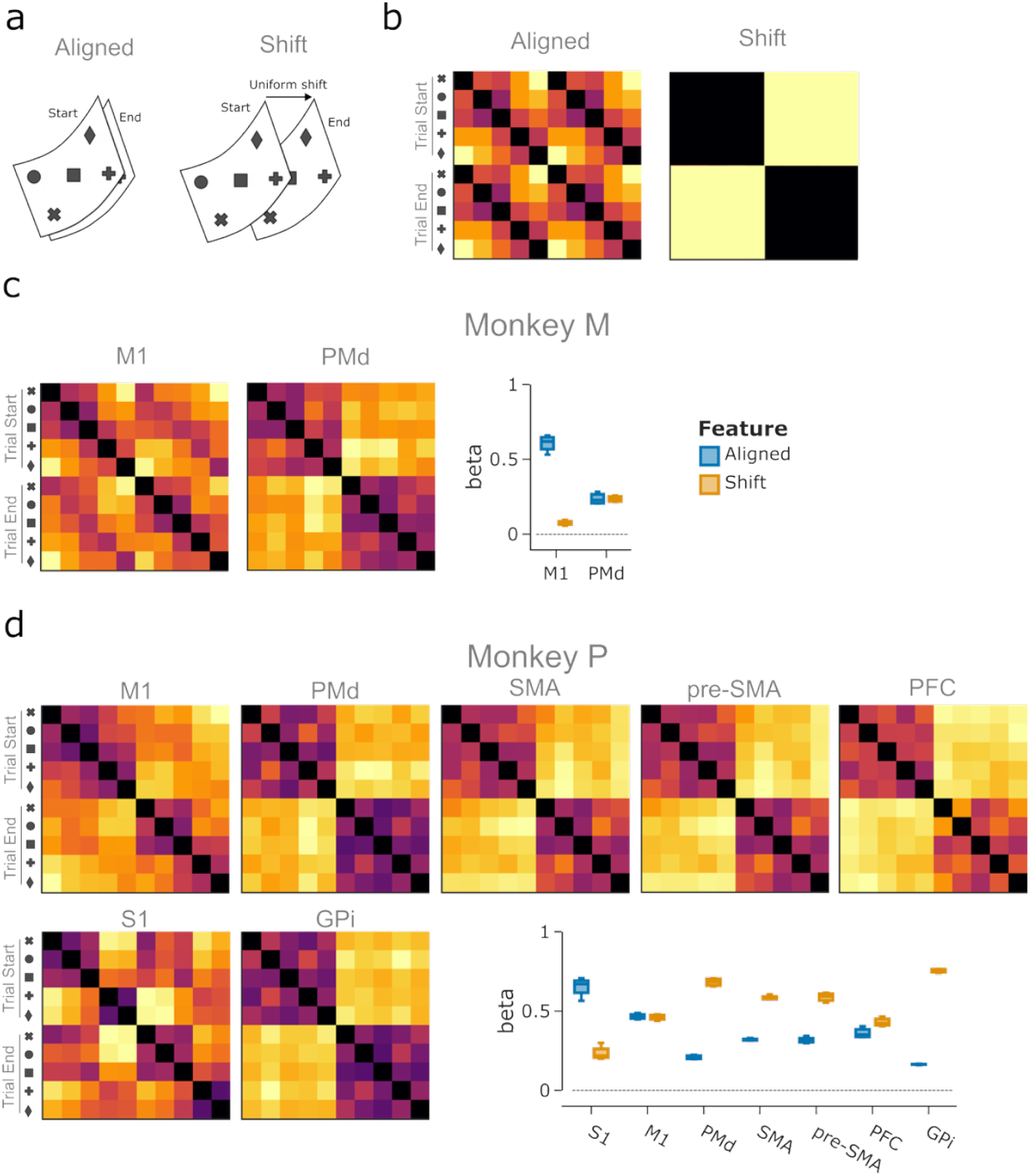
Geometry of neural state space at start and end posture. **a)** Schematic illustration of the *Aligned*, and *Shift* models. In both, the within-Start and within-End fixed point structures are identical. In the *Aligned* model, fixed points at the Start and End of the trial fully overlap, whereas in the *Shift* model, Start and End fixed points are separated by a uniform, condition-independent shift in neural state space. **b)** Representational dissimilarity matrices (RDMs) corresponding to the *Aligned* and *Shift* models. **c)** RDMs derived from neural activity when the monkey*’*s hand was stationary at each of the five target locations, either at the Start or End of the trial, shown for M1 and PMd in Monkey M. Box plots show non-negative regression beta weights quantifying the contribution of each model to the observed neural geometry. **d)** Same as (c), but for all recorded brain regions in Monkey P.

To test these predictions, we constructed RDMs from the full-dimensional neural activity for each target at trial start and end (**Fig. 5c**, e.g. M1). Consistent with the first prediction, the geometry across the five fixed points was preserved: distances within each time point were very similar (**Fig. 5c, M1**, diagonal blocks). However, comparing across start and end points revealed a near-constant offset (**Fig. 5c, M1**, off-diagonal blocks), suggesting a uniform shift of the fixed points in state space. We quantified uniformity of this shift by computing vector differences between start and end fixed points for random condition pairs and measuring their alignment (dot product). The shift vectors were consistently parallel (**Supplementary Fig. 8a**), confirming its uniformity. To quantify the contribution of aligned geometry and uniform shift, we defined a Shift component (**Fig. 5a–b**, left) that assumes identical geometry at both time points but a constant offset between them. Using non-negative linear regression, we fit both the Aligned and Shift geometries to the data (**see Methods**).

In monkey M, the Aligned component showed the largest contribution in M1, with only a small contribution from the Shift. In PMd, both models contributed equally. In monkey P, only S1 and M1 showed a dominant contribution from the Aligned component; all other areas—including SMA, pre-SMA, PFC, and GPi—showed larger contributions of the Shift. Nonetheless, in all areas and both monkeys, the Aligned component had a beta weight significantly above zero, suggesting that alignment is present in all regions, even if it contributes less in some. To reconfirm that the aligned geometry persists despite the uniform shift, we trained a classifier to decode resting hand location using the neural data at the start of the trial and tested it on the same targets at the end of the trial. Classification accuracy remained well above chance in all regions, reinforcing the idea that the same fixed point geometry persists, despite the presence of a condition-independent shift (**Supplementary Fig. 8b**).

Together, these results reveal that all areas show a condition independent, uniform shift in neural activity over the trial duration, superimposed on a stable fixed point structure. This uniform shift was not visible in the original PCA projection as the PCs are optimized to highlight differences across conditions, rather than components shared across them.

### Dynamics of the uniform shift dimension

Comparing neural states at rest before and after movement revealed a uniform shift in neural state space. Because the arrangement of posture fixed points remains consistent at the start of every trial, we reasoned that this shift must fully reverse during the interval between trials. To characterize the full trajectory of this return, we projected neural activity from both the trial and the between-trial interval onto the uniform shift dimension—defined as the axis of maximal difference between the pre- and post-movement rest periods (see Methods). At the end of each successful trial, following reward delivery and a fixed delay period, the appearance of a new home target prompted the monkeys to move to the next home target and initiate the next trial. While this between-trial movement was unrewarded and less controlled, it provided a critical window to observe the neural reset. We therefore aligned both kinematic and neural data to the onset of this new home target cue (**Fig. 6**) to track the state space dynamics as they returned to baseline.

**Figure 6.**
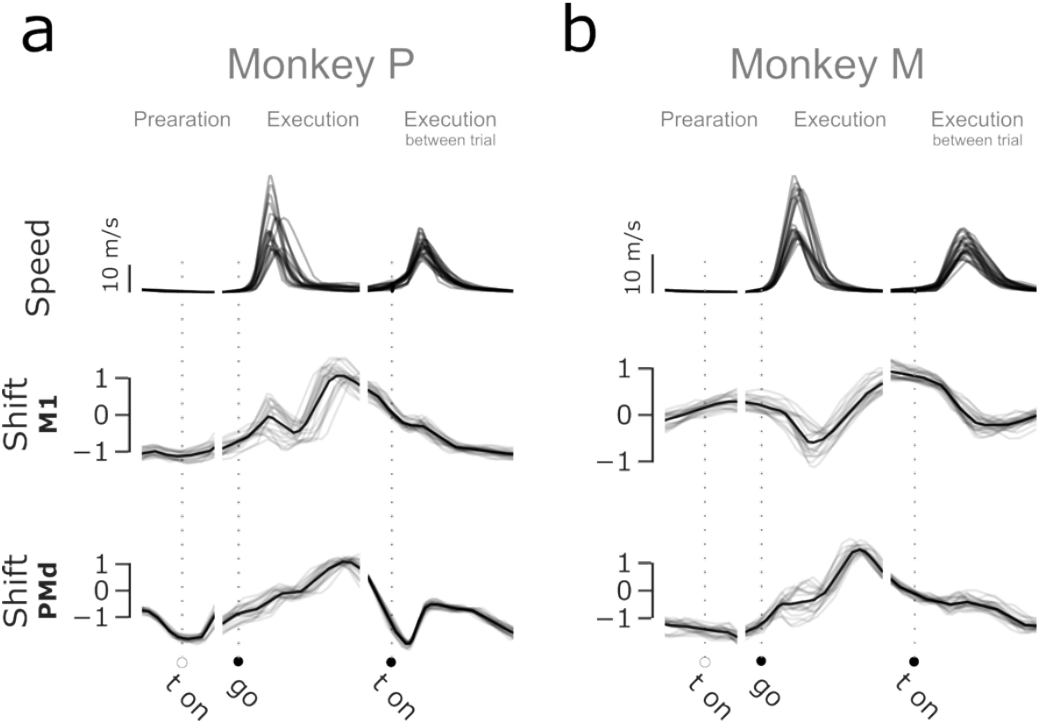
Projection of neural data on the uniform shift dimension. **a)** Speed profile (top), and projection of M1 (middle) and PMd (bottom) neural data onto the uniform shift dimension during the within-trial preparation and execution phases, as well as the between-trial movement. The withing trial preparation and execution phases are aligned to target onset and the go cue respectively. The between-trial phase is aligned to the subsequent home target onset. **b)** Same as (a) but for Monkey M.

In Monkey P, for both M1 and PMd, most of the change in the shift dimension occurred during the trial*’*s movement phase, peaking once the hand reached the final target (**Fig. 6a**). Notably, the projection returned fully to baseline during the between trial phase as the monkey moved toward the next home target; we observed the same pattern in Monkey M (**Fig. 6b**).

The activity within the uniform shift dimension occurred over a significantly longer timescale than that of typical single-movement execution. This suggests that the uniform shift axis is functionally distinct from the well-studied *condition-independent signal (CIS)*^45^ components associated with movement production. To directly compare these signals, we projected the neural data onto the CIS dimensions optimized during the execution phase (**Supplementary Fig. 9**; See Methods). For both M1 and PMd, the temporal occupancy of these dimensions differed: while CIS dimensions were reliably activated during both the trial movement and the inter-trial movement, activity within the shift dimension slowly ramped up during the main trial and ramped down during the inter-trial phase (**Supplementary Fig. 9**). This distinct temporal profile suggests a unique functional role for the uniform shift in monitoring high-level task information—such as trial progression—alongside the shorter-timescale signals that govern movement execution.

### Posture-dependent control shapes neural geometry

Our results demonstrate that motor cortical areas preserve a representation of posture while simultaneously generating the dynamics that drive muscle activity and exhibit a contextual shift that distinguishes movement onset from termination (**Fig. 3**). The geometry of neural dynamics is organized such that posture dimensions, movement control dynamics, and context-dependent shifts are all continuously updated throughout movement execution. Why do cortical areas retain both location and progression information during movements? One possibility is that such a representation is essential for controlling effectors with complex dynamics, where the control policy depends on current position^19,20^—as in the arm.

To test this idea, we trained recurrent neural networks (**Fig. 7a**) on the same delayed-reach task but with effectors of varying complexity: a point-mass model, in which the control policy is independent of position, and a two-joint, six-muscle arm with intersegmental dynamics (**Fig. 7b**). Networks received the target*’*s XY location, and a go cue as input, and output muscle activity or force commands to the plant. The architecture comprised three modules: a PMd-like recurrent network receiving task inputs and visual feedback; an M1-like module connected to PMd-like module and sparsely to the task and visual signals; and a spinal cord–like module receiving proprioceptive feedback and generating muscle activity. Plant complexity was the only difference between the two networks; connectivity, architecture, and training procedures were identical. Seven independent networks were trained for each effector type (**Fig. 7a**, see Methods).

**Figure 7.**
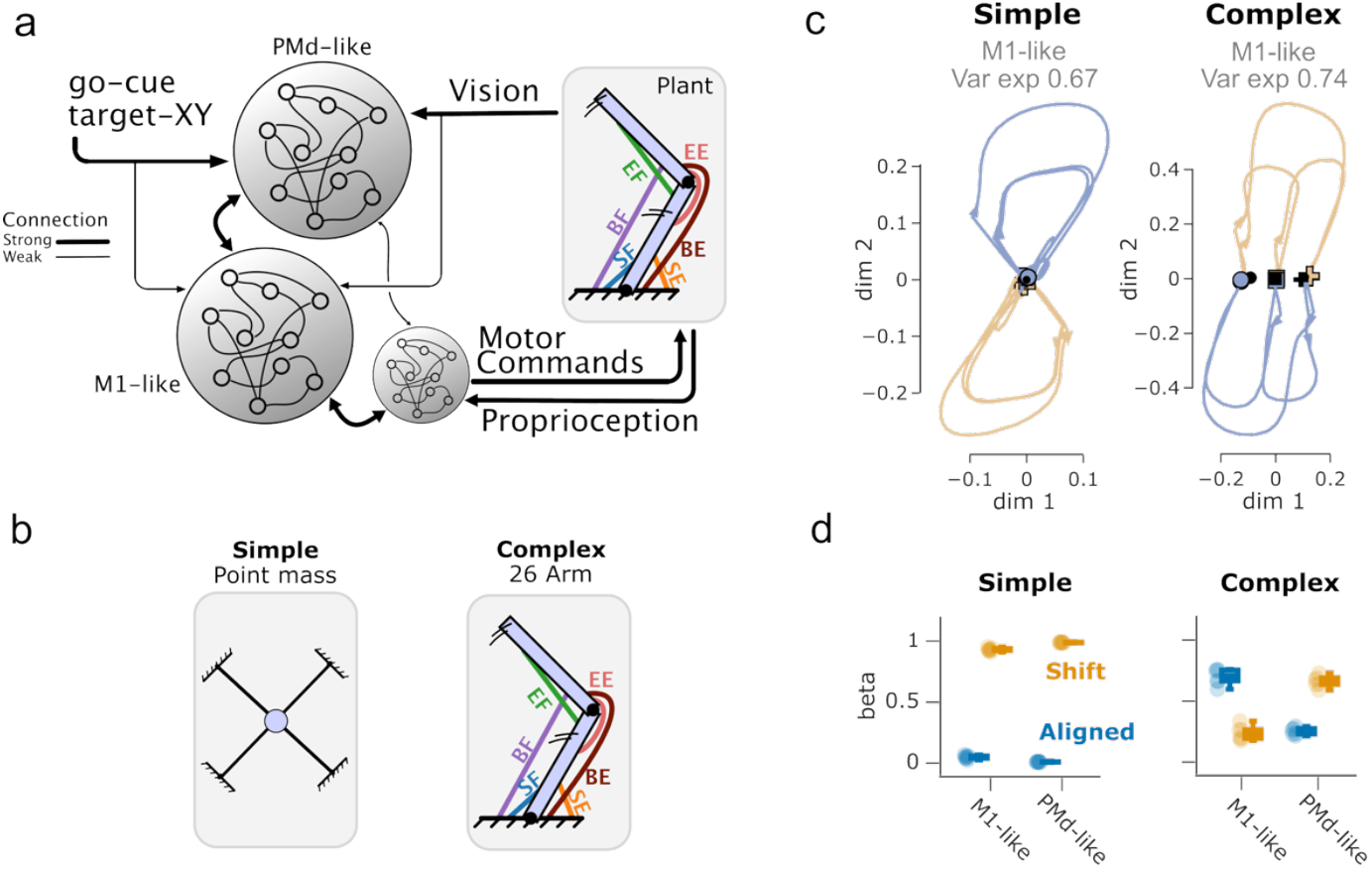
Effector complexity shapes the geometry of neural dynamics. **a)** A three-module network architecture. Arrows show the connections and their strength (see Methods). **b)** Effectors of varying degree of complexity. Simple effector consists of a points mass with four linear actuators. Complex effector with two joints and six muscles. **c)** Neural trajectories for M1-like module. Like figure 3b. Dotted traces represent center-out reaches. Each trace begins at a black marker indicating the start location, shaped as a plus, square, or circle corresponding to trials that began from the plus, square, or circle target, respectively. Traces are colored by reach direction, and arrows indicate the go cue. **d)** Shift and Aligned contributions. Like figure 6C. Box plots show non-negative regression beta weights quantifying the contribution of Shift and Aligned models to the observed neural geometry.

Both networks achieved high accuracy. However, consistent with our prediction, posture representation during movement was markedly reduced in networks controlling the point mass (**Fig. 7c**, simple) but robust in those controlling the complex arm, which reproduced five posture-specific fixed points, similar to that observed in the neural data (**Fig. 7c**, complex). Quantifying PMd-like and M1-like geometry revealed that simple-effectors produced similar alignment and shift values in both modules (**Fig. 7d**, simple), whereas complex-effectors showed clear differentiation: M1-like modules exhibited strong alignment components, while PMd-like modules were dominated by the shift component—closely matching the organization seen in monkey M1 and PMd (**Fig. 7d, Fig. 5c**).

These simulations provide a normative explanation for the geometry observed in the data: posture-specific structure emerges naturally in networks controlling effectors whose dynamics require location-dependent control policies.

## Discussion

We designed a task in which each target location served both as a start point on some trials and an end point on others, allowing us to examine neural dynamics across a variety of postures. This approach revealed compositional structure in motor cortical activity comprising three key components. First, a posture subspace containing fixed points for each target, visited whenever the arm rests at that location before or after a reach. Second, rotational dynamics linking these posture fixed points, arranged such that similar rotations correspond to reaches in similar directions. Importantly, the projection of these rotational dynamics onto the posture subspace provides continuous updates on limb posture. Third, superimposed on the other components, a uniform shift dimension reflecting trial progression. This structure reveals how motor cortical areas can simultaneously generate the activity required for movement, continuously update the effector*’*s current location, and track progress within the trial.

In the postural subspace we found that neural states corresponding to different postures were organized not only by the physical distances between targets in the workspace, but also according to an equidistant, orthogonal geometry (**Fig. 3-4**). As expected, physical distance coding (in cartesian or joint space, **Supplementary Fig. 7**) was strongest in S1 and M1—areas most directly connected to the brain stem and spinal cord. However, we were somewhat surprised to observe any abstract coding in M1. The presence of more abstract coding in anterior regions was less surprising, given prior evidence of their involvement in higher-order functions such as decision-making^10,46,47^. One important caveat is that our task involved only five fixed targets, and the monkey was highly trained to reach between them. This may have inflated the prominence of abstract representations for these specific locations, leaving open the question of whether similar coding would emerge in a larger, more varied target set, or how quickly it would develop during training. Nevertheless, associating final goals with an abstract neural code—even in motor areas like M1 and PMd—could provide a powerful mechanism for coupling novel control dynamics for moving to the same final goal, as it is useful for learning to adapt to external perturbations^48–51^.

We found that the neural dynamics underlying both movement preparation and execution were primarily organized by movement similarity (**Fig. 3, Supplementary Fig. 5**). In our experiment, we manipulated movement start location, direction and extent. Among these variables, direction was most important in explaining the geometry of neural trajectories. This is expected as movements with different directions demand significantly different muscle activity patterns. Movement extent was not independently encoded but rather appeared only to modulate the direction code (**Supplementary Fig. 5**). Thus, our results suggest that movement between two points is controlled by combining the neural state for the starting posture with a movement plan, as previously suggested by Marino et. al.^35^. However, in addition, our results also suggest that this movement plan encodes direction and extent jointly, rather than in separate, orthogonal dimensions (**Supplementary Fig. 5**). In the simulations we also showed that complexity of effector can explain the arrangement of neural dynamics in the state space: more complex effectors requiring posture-dependent control policies demand a more robust representation of posture within the neural state space (**Fig. 7**). Taken together, our data and simulations suggest that the organization of neural dynamics reflects the network-level strategies necessary for generating complex muscle activity patterns^52^.

With our task design, we were able to compare the neural states associated with the same postures at both the start and end of a reaching movement. This revealed a distinct neural representational geometry, characterized by two components: First, we found an *aligned* component, in which the dynamics fully return to the same place for the same posture regardless of whether it occurs at the start or end of a trial. Additionally, we found a *shift component*, which is similar for all postures and only separates start and end states of the trial (**Fig. 5**). The alignment component was strongest in M1 and S1, consistent with their relatively direct roles in controlling posture and processing sensory feedback via spinal circuits^53,54^. Conversely, the shift component was more prominent in other cortical areas, suggesting functional specialization across the motor hierarchy. In line with this, we observed the same pattern in recurrent neural network simulations only when the network was trained to control a complex effector. Interestingly, the shift component was *condition-independent*: it appeared similar regardless of which start and end postures were compared. This raises the question of how it relates to the previously reported condition-independent (CI) dimensions that occur at the transition between posture and movement^45^. The previously reported CI dimension is defined in center-out tasks as the dimension that captures the largest variance around movement onset and is similar across reach conditions. While the shift component we report shares the property of being similar across conditions, it differs in an important way: in the previously reported CI dimension, activity peaks around movement onset and then returns to baseline by the end of the reach. In contrast, in areas with a strong shift component, the activity along the shift dimension increases gradually and does *not* return to baseline at movement end (**Supplementary Fig. 9**). What might be the benefit of having such a shift dimension? One possibility is that it provides a representation of task progression. However, It does not appear to be an absolute timing signal^55^ since progression along the shift dimension was similar for faster (long extent) movements.

The geometry of the neural dynamics we observed in motor areas were highly consistent across recording sessions, even though the individual neurons recorded were different from session to session. This is consistent with previous work showing that neural geometries can be aligned with high accuracy not only across sessions^56^ but even across animals^57^. Returning to the same fixed point via specific rotational dynamics was similar to the results of Oby et al., who used a brain–computer interface (BCI) to allow a monkey to control a cursor*’*s position via primary motor cortex activity^58^. In their study, motor cortical dynamics also moved between two fixed points associated with instructed cursor locations. Notably, they found that it was nearly impossible to deviate from these neural trajectories during BCI control, suggesting it they are largely dictated by local connectivity rather than by external inputs. Our results indicate that these patterns are not special to BCI control, rather it suggests that postural equilibrium states^59–64^ may constitute a key organizing feature of motor cortical dynamics.

Taken together, our results offer the following picture of motor cortical dynamics during reaching (**Fig. 8**). At rest, the neural state is confined to a posture manifold^65,66^ that continuously represents limb state. When movement to another posture is required, the neural state evolves orthogonal to the posture^35^ manifold—this constitutes motor preparation^8^. At the go cue, specific rotational dynamics^7^ move the neural state back to the posture manifold at the desired final posture (**Fig. 3**). These rotational dynamics, when projected onto readout neurons in the spinal cord, generate the required muscle activity^3^. Rotational dynamics are organized such that more similar rotational dynamics lead to more similar movements (**Fig. 5**). In higher-level motor cortical areas this dynamical structure is superimposed on a uniform shift dimension that evolves during movement execution and allows other readout neurons to directly track movement progress (**Fig. 8**, PMd-like neural state). While this work provides the most comprehensive account of motor control dynamics during reaching to date, additional work is needed to determine how other movement parameters, such as movement speed^52^ or curvature^67,68^ are encoded, and how these representations change with learning^48,69^.

**Figure 8.**
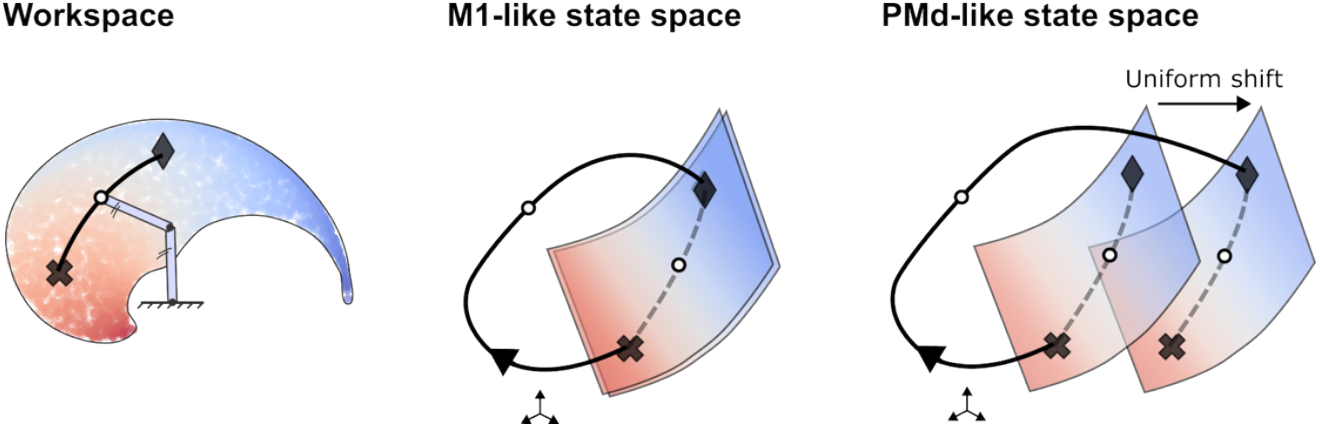
A basic sketch for the geometry of motor cortical neural dynamics. **A**) Two-dimensional workspace for a two-joint arm. Colors indicate joint states from fully flexed (red) to fully extended (blue). Sample reach between two points (cross and diamond) with hand path (black trace) and current endpoint location (white circle). **B)** Neural trajectory associated with the example reach in an M1-like state space. The posture manifold is colored by workspace locations. Black trace shows neural trajectory for the sample reach; white circle indicates current endpoint location; dotted line shows trajectory projection onto the posture manifold. **C**) Same as but for PM-like state space, where a uniform shift is superimposed with the posture manifold, separating states associated with trial start and end.

## Methods

### Subjects

Two male rhesus macaques (Macaca mulatta) participated in this study: Monkey M (16 years old, 10 kg) and Monkey P (11 years old, 16 kg). All procedures were approved by the Institutional Animal Care and Use Committee at Western University (Protocol #2022-028).

### Apparatus

Experiments were conducted using a KINARM^38^ exoskeleton robot. Monkeys preformed the task with their right arm positioned in the exoskeleton, which supported the arm*’*s weight and allowed free movement in a horizontal plane. A circular cursor (0.5 mm diameter) representing the position of the index fingertip, was displayed on a horizontal screen that occluded direction vision of the arm (**Fig. 1a**). Hand kinematics were recorded at 1000 Hz.

### Task

Each trial began by presenting one of the five possible targets as a start home target. The potential target locations were positioned at four corners and center of a 10 × 6 cm rectangle (**Fig. 1b**). After the monkey held his hand withing the start target for 500 ms, one of the four remaining targets was randomly selected and displayed as the reach end target. The monkey was required to maintain his hand withing the start target for a variable delay period of 300-700 ms. The go cue was then given by extinguishing the start target and playing an auditory tone (800 Hz). The monkey initiated the reaches promptly and received a juice reward upon maintain the hand withing the goal target for 150 ms (**Fig. 1c**). The next trial began after one second of iter-trial interval. Reach conditions were defined by all possible combinations of start and end targets, resulting in 20 unique conditions. These conditions were fully randomized withing each session.

### Neural recordings

The setup and procedures used here were similar to our previous work^14^. Both monkeys were implanted with a custom 3D-printed titanium head cap that included an integrated recording chamber and head post. For each animal, the implant was individually designed to conform to the skull surface based on a micro-CT scan. To identify recording targets, CT images were registered to a high-resolution structural MRI acquired prior to surgery. Following alignment, we warped anatomical segmentation from a macaque brain atlas (CHARM^70,71^ for cortical areas and SARM^72^ for subcortical areas) to each monkey*’*s individual MRI. This allowed us to precisely plan recording trajectories relative to the implant geometry. In monkey M, we validated the accuracy of our targeting post-mortem and confirmed that planned trajectories reached cordial targets with <0.5 mm deviation. A large craniotomy was performed in Monkey M to access all planned recording sites, whereas in Monkey P, five 2.7 mm burr holes were drilled at the desired entry points. Recording in both animals were performed using Neuropixels 1.0 NHP Long probes^73^. Probes were mounted in a custom-designed holder equipped with a 9 mm retractable guide tube. For each recording session, we 3D-printed an adapter (Formslab 3B+, Gray Resin V4) that attached the probe holder to the titanium recording chamber, aligning the probe along a pre-planned trajectory. In Monkey M, the printed adapter (Formslab 3B+, BioMed Clean Resin) included a stabilizing extension that rested on a region of the craniotomy not used for the recording to improve mechanical stability.

### Neural data preprocessing

We used SpikeGLX for neural recordings and performed spike sorting using a custom pipeline (https://github.com/mehrdadkashefi/pixelpipe). Preprocessing steps included: applying a high-pass filter with a 400 Hz cutoff frequency, removing noisy channels, correcting small phase shift between the recorded channels, and applying a common average referencing filter. These steps were implemented in SpikeInterface^74^. Following these steps, data were spike-sorted using Kilosort 4^75,76^ with its built-in drift correction. Across all recordings, we observed minor drift (< 50 um), which was corrected by Kilosort*’*s drift correction. A List of recording session along with number of successful trials and recorded neurons in each session can be found in Supplementary table. Single units were included for analysis if they met the following criteria: (1) labeled as *‘*good*’* by Kilosort using the default parameters. (2) had an average firing rate above 0.1 Hz across the recording session, and (3) demonstrated temporal stability. Stability was assess using a modified relative standard deviation metric. We divided each recording session into ten equal partitions and computed the mean firing rate of each neuron within each partition. A single neuron was considered stable if the standard deviation of its firing rate across partitions was less than 30% of its overall mean firing rate. This approach excluded neurons that were inactive or exhibited large fluctuations during parts of the recording session. The number of remaining high-quality isolated neurons is reported for each area in **Fig. 2b** and **Supplementary Fig. 1**. Following spike sorting, spiking activity for each single unit was aligned to both target onset (-500 ms to +400 ms) and go cue (-200 ms to 1400 ms) on a per-trial basis. Spike trains were then smoothed using a causal Gaussian kernel with a 25 ms standard deviation to obtain a trial-by-trial estimate of firing rate. These trial-level firing rates were subsequently averaged across trials for each condition to obtain a denoised estimate of the firing rate, resulting in a data tensor of shape condition x time x neurons for each session. For every condition, at least 25 trials were averaged per session. After alignment and averaging, neurons recorded from the same brain area were concatenated across sessions. All analysis were performed at a 1000 Hz sampling rate.

### Single neuron tuning analysis

For the analyses in **Fig. 2b, Fig. 2c**, and **Supplementary Fig. 1**, we used a classification-based approach to determine whether individual neurons were tuned to start position, movement direction, or extent. This analysis was performed separately for the preparation and execution phases. For each recording session, we constructed a trial × time × neuron data matrix aligned to the go cue. To assess tuning during the preparation phase, we focused on the window from 550 ms to 50 ms before the go cue. Firing rates were averaged within this time window for each trial, resulting in a trial × neuron data. We then used logistic regression^77^ (Scikit-learn^78^ implementation, with no penalty) to classify start position at the single-trial level based on the activity of individual neurons. Classification performance was evaluated using 10 × 10-fold cross-validation. To establish a significance threshold, we repeated the classification procedure using shuffled condition labels to generate a chance-level distribution. A neuron was considered tuned to a given parameter if its classification accuracy exceeded the accuracy of the shuffled-label control. To assess tuning for movement direction, we used the same classification approach described above. However, because start position can partially predict movement direction, we performed separate classifications within each start position. This ensured that, within each subset, all possible directions were equally likely. A neuron was considered direction-tuned if it showed significant decoding performance for any of the five possible directions from at least one start position. To evaluate tuning to movement extent, we restricted the classification to conditions in which extent varied while start position and movement direction remained constant—specifically, the diagonal reaches. This controlled for potential confounds between extent and other movement parameters. The same analysis procedure was repeated for the execution phase, using neural activity averaged from 50 ms to 550 ms after the go cue. Once tuning was determined for each neuron with respect to start position, direction, and extent, we grouped neurons into categories based on the number and combination of parameters to which they were tuned: position only (P), direction only (D), extent only (E), pairs of parameters (PD, DE, or EP), or all three (PDE). Neurons that were not significantly tuned to any parameter were categorized as *“*untuned.*”* The proportions of neurons in each category were summarized in the pie charts (**Fig. 2b–c**). We applied the same classification approach for **Supplementary Fig. 2**, but instead of averaging firing rates over a time window, we performed the regression at each time point. The onset of significant decoding (indicated by a red line in Supplementary Fig. 2) was defined as the first time point at which classification accuracy exceeded chance and remained above chance for at least 100 ms.

### PCA analysis

For **Fig. 3** and **Supplementary Fig. 6**, we used principal component analysis (PCA) to visualize the geometry of neural population activity. Neural data were aligned to the go cue (–400 ms to +1400 ms) and trial-averaged within each condition, resulting in a condition × time × neuron tensor. For each brain area, neurons recorded across different sessions were concatenated. Prior to PCA, we centered and soft-normalized the firing rates by subtracting each neuron*’*s mean firing rate and dividing by its dynamic range (i.e., maximum – minimum firing rate) plus a constant offset of 5 spikes per second^7,18,79^. This soft normalization ensured that neurons with large firing variability did not dominate the principal components, allowing more uniform contribution across the population. We then reshaped the condition × time × neuron tensor into a two-dimensional matrix of size (conditions × time) × neurons and applied PCA. We then plotted the data projected on either top two three PCs. For each area, the variance explained by these top PCs are shown in **Supplementary Fig. 6**. Within the PC space the orientation of data can be different for different PCA fits. To ensure that the posture axis always aligns with x-axis, we performed a second PCA with the same number of dimensions just to rotate the data. This second PCA was fit on the data 50 ms before target onset to find the rotation matrix that aligns spread of the posture fixed points during that period onto x-axis. Note that this PCA step does not change the amount of variance explained and it is just performed have better visual presentation of the data. We then visualized the neural trajectories by projecting the data onto the top two or three principal components. For each brain area, the variance explained by these components is shown in **Supplementary Fig. 6**. To facilitate consistent visualization—specifically, to align the posture-related variance with the x-axis—we applied a secondary PCA (with the same number of dimensions) solely for rotation purposes. This second PCA was fit on neural activity from the 50 ms preceding target onset, and its principal axis was used to rotate the entire dataset so that the spread of postural states during this period aligned with the x-axis. Importantly, this reorientation did not affect the amount of variance explained or the underlying geometry; it was applied purely to enhance the visual interpretability of the plots.

### Isolated posture, preparation and execution subspaces

To isolate the subspaces associated with posture, preparation, and execution, we followed previous optimization-based approaches^9,16,18,36^ to identify three mutually orthogonal subspaces. These subspaces were designed to explain the maximum across-condition variance for three distinct task epochs: posture, preparation, and execution. For the *posture subspace*, we utilized a 500 ms window prior to target onset (*T*_*post*_), during which the monkeys*’* hands remained stationary within the home target; this period captures neural activity purely associated with posture across all conditions. For the *preparation subspace*, we used a 500 ms window following target onset but prior to movement initiation (*T*_*prep*_), which contains both postural and planning information. Finally, for the *execution subspace*, we defined a 1000 ms window immediately following the go cue to capture movement-related activity(*T*_*exe*_). All preprocessing steps (including range normalization and removal of the cross-condition mean) were consistent with the PCA procedures described above. We then sought three sets of dimensions: posture (*W*_*post*_), preparation (*W*_*perp*_), and execution (*W*_*exe*_). We desire these subspaces to be mutually orthogonal while maximizing the captured variance for their respective epochs 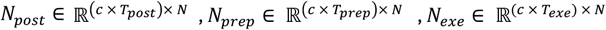. To achieve this, we solved the following optimization problem:

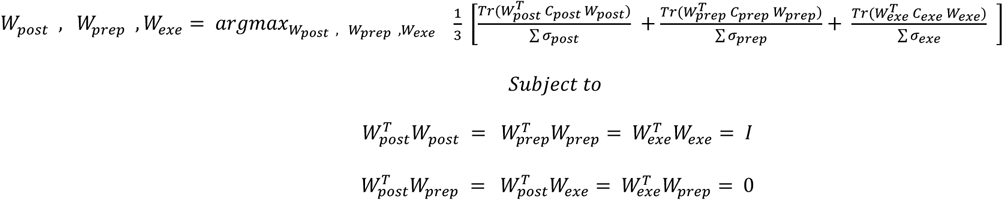

Here *C*_*post*_, *C*_*perp*_, *C*_*ext*_ are covariance matrix for neural data at each respective epoch. *σ*_*post*_, *σ*_*perp*_, *σ*_*exe*_ are singular values associated with each covariance matrix. Tr(.) denote the trace operator, hence the numerator in the loss function, 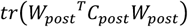, yields the total variance in the posture data explained by the optimized posture subspaces. These values are normalized by the sum of associated singular values, so the optimizer is insensitive to differences between total variance in different epochs. We use Pymanopt^80^ with Stiefel manifold for solving the optimization problem. With 20 dimensions for each of the subspaces, we were able to explain able to explain ∼75 percent of variance for in posture, preparation, and execution phases. Note that this optimization process does not return sorted dimensions. Hence, we applied a secondary PCA (with the same number of dimensions, hence with no loss of variance) on the optimized Ws. This step ensures that the optimized subspaces are order such the first one captures most of variance and so forth. This step was particularly useful for (**Supplementary Fig. 3**) in which we only show top two dimensions of each subspace.

### Quantifying strength of rotation dynamics

Simple PCA indicated that during execution, the neural state transitions between postural fixed points with rotational dynamics. To directly quantify the strength of these rotations, we applied *jPCA*^7^ to the execution-phase activity (**Supplementary Fig. 4**). We first aligned the neural data to the go cue (spanning -200 ms to +1000 ms) and projected it onto the three previously optimized subspaces for posture, preparation, and execution. This resulted in a 20-dimension x Time x Condition dataset for each subspace, capturing the evolution of neural activity during movement with each subspace. For each projection, we fit a linear dynamical system (**Supplementary Fig. 4a)** to predict the state at the next time point using the following equation:

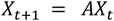

Following the jPCA framework, we performed two separate fits: one where the transition matrix *A* was unconstrained (free matrix) and another where *A* was constrained to be *skew-symmetric*. A skew-symmetric matrix *A* yields purely imaginary eigenvalues, representing perfectly rotational dynamics. We then quantified the strength of the rotations by comparing the variance explained by the skew-symmetric model against the unconstrained linear model. This analysis was conducted independently for the execution-phase data projected onto the posture, preparation, and execution subspaces (**Supplementary Fig. 4b-c**).

### Projection on condition-independent and shift dimensions

In **Figure 6** and **Supplementary Figure 9**, we projected neural activity onto condition-independent and shift dimensions. The condition-independent signals (CIS) are subspaces that are invariant to experimental conditions, instead they maximize variance related to time. To extract these, we utilized demixed principal component analysis (dPCA^81^), fitting the model using neural data aligned to the go cue (from -300 ms to +700 ms). Briefly, dPCA assigns labels to each data point indicating its condition or time identity, then optimizes a set of dimensions (*K*) that maximize variance related to condition, time, or their interaction. For this analysis, we specifically utilized the *K* dimensions that covary with time as our condition-independent components.

To estimate the shift dimension, we identified the direction that maximizes the difference in neural activity between postural states at the beginning versus the end of the trial. This was calculated by subtracting the mean neural activity during the final 200 ms before target capture (end of trial) from the mean activity during the 300 ms prior to target onset (start of trial). In both windows, the monkeys*’* hands remained stationary across the five targets, ensuring the shift dimension specifically captured changes in baseline postural activity.

### Analysis for quantifying neural geometry

#### General framework

For **Figures 4–5 and Supplementary Fig. 5**, our primary goal was twofold: (1) to summarize and visualize the geometry of neural population activity, and (2) to quantify how well this data geometry matches a set of hypothesized geometries. To accomplish this, we used a representational model framework^82^, in which the key statistical quantity is the second moment matrix of the activity patterns. Let *U* be a (*K × P)* matrix representing the true firing rates of all recorded neurons (*P*) across (*K)* experimental conditions at a given time point. The (*K × K)* second moment matrix *G* is then computed as:

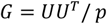

This matrix captures the variance of neural activity for each condition (on the diagonal) and the covariance between conditions (on the off diagonal). To summarize and visualize the geometry, we plotted the representational dissimilarity matrix (RDM), which contains the pairwise Euclidean distances between condition-specific activity patterns (rows of *U*). The Euclidean distance between conditions *i* and *j* can be computed directly from *G*:

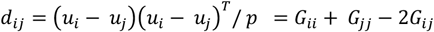

To model the compositional structure of the neural geometry, we described the empirical second-moment matrix as linear combination of different model components. This is equivalent to the assumption that the different components are represented in orthogonal subspace of the neural code^82^. We estimated the contribution of each component by performing a non-negative linear regression to explain the empirical second moment matrix obtained from data *G*_*emp*_ as a weighted combination of component matrices:

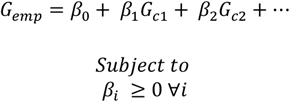

Before regression, all *G* matrices (both data and components) were normalized by dividing by their trace^83^, ensuring that each component contributed to a unit total variance. This normalization allows the resulting β coefficients to be interpreted as the relative contributions of each component to the observed second moment.

In all the analysis, for brain areas with multiple recording sessions, we estimated the data second moment matrix *G*_*emp*_ and corresponding β coefficients separately for each session and reported the resulting β distributions in box plots. For brain areas with only a single recording session, we performed 10 repetitions of 5-fold cross-validation across trials within that session. In each fold, we computed the condition-averaged firing rates, constructed the *G*_*emp*_, and estimated the β coefficients—repeating the full procedure 10 times. Across sessions—despite small differences in penetration locations and the number of neurons recorded—we observed strong consistency in the structure of the estimated *G* matrices. Specifically, the average cosine similarity between *G* matrices across sessions exceeded 0.9, indicating highly reliable covariance structure in the population activity^56,57,65^. Because of this robustness, we did not apply cross-validation when estimating the second moment matrix which is common practice in modalities with lower signal-to-noise such as fMRI^84,85^. For components that involved parameter fitting (e.g., DE Orthogonal, **Supplementary Fig. 5**, or shift + alignment model, **Supplementary Fig. 5**), we ensured proper model validation by tuning parameters only on the training session or fold and evaluating performance on held-out data.

#### Geometry at start posture

For the analyses related to posture at the start of the trial, none of the model components involved any trainable parameters. The *G* matrix for the *Identity* component was simply a 5 × 5 identity matrix. The *Cartesian* component was derived directly from a centered 5 × 2 feature matrix **F**, containing the (x, y) coordinates of the five target locations in the workspace. The component *G* matrix was computed as:

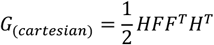

where *H* is a 5 × 5 centering matrix.

To estimate the data *G* matrix for each recording session, we first computed the average firing rates for all conditions sharing the same start location, within a 150 to 50 ms window before target onset (when the monkey*’*s hand was stationary). This yielded a 5 × *P* matrix *D*, where *P* is the number of recorded neurons in the session. We then centered *D* and computed the empirical *G* matrix as:

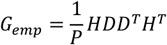

#### Geometry at peak speed

The empirical *G* matrix was estimated as described above, using 100 ms of neural activity centered on the time of peak movement speed. Peak speed was determined from the average speed profile for each of the 20 experimental conditions. This analysis evaluated four hypothetical geometries of neural state space. The *Direction-only* model component (**Supplementary Fig. 5a**, Direction) assumed a ring geometry representing all possible reach directions. Such a geometry is expected when neurons exhibit smooth directional tuning^86^ (e.g., cosine tuning). We simulated a ring with points for each of the experimental direction and generated a feature matrix *F* (conditions × 2), where each row corresponded to the coordinates of the direction on the ring associated with one experimental condition. The model *G* matrix was then computed from *F* using the same procedure as above. The *Extent-only* model component (**Supplementary Fig. 5a**, Extent) encoded movement extent as a linear variable. The corresponding feature matrix contained a single value per condition, representing the reach extent. Both the Direction-only and Extent-only model components had no trainable parameters. The *Direction-Extent (DE)* model component assumed a geometry of three concentric rings, one for each movement extent. Direction was coded by the angular position on each ring, while extent was coded by the ring*’*s radius. This geometry would arise if neurons are tuned for direction with a smooth function and modulate their firing rate with movement extent. The relative radii of the rings were scaled to match the ratio of movement extents in the experiment. The *Direction-Extent Orthogonal (DE Orthogonal)* model component represented direction and extent in orthogonal dimensions (**Supplementary Fig. 5a**, DE Orthogonal). Here, three rings (one per extent) were arranged in parallel planes, with the inter-ring separation in the orthogonal dimension encoding extent. Both the ring radii and the separation between rings were treated as tunable parameters, optimized on the training data. For both DE and DE Orthogonal model components, we constructed a 20 × 3 feature matrix *F* by selecting points in the model geometry that matched the direction and extent of each experimental condition. The corresponding *G* matrices were then calculated as before.

#### Geometry at Start vs End of trial

In this analysis, we aimed to compare the neural states associated with the five posture locations—whether the arm was resting at those locations at the *start* or the *end* of the trial. To estimate the neural state at the start of the trial, we used a 100 ms window from 150 to 50 ms before target onset. To estimate the neural state at the end of the trial, we used a 200 ms window immediately before the end of the trial. In both time windows, the arm was stationary at the target locations. We then averaged neural activity across time points and across all trials sharing the same posture (start or end), resulting in two matrices of size 5 × *P* (where *P* is the number of recorded neurons)—one for the start and one for the end. These were concatenated into a single data matrix *D* of size (10 × *P)*. The empirical *G* matrix was then computed from *D* using the same procedure described earlier. This resulted in a (10 × *10)* empirical *G* matrix that captures the geometry of posture location at both Start and End of the trial.

To construct the model component, we first considered a *fully aligned* component. This model component assumes that the neural states associated with each posture are identical at the start and end of the trial. To build it, we estimated the covariance structure of the five posture locations using neural activity from the start of the trial (on held-out data) and assumed that the same structure would apply to the end of the trial as well as to comparisons between start and end. As a result, the model component *G* matrix consisted of four identical blocks: one for within-start, one for within-end, and two for start–end cross terms (Fig. 5B, Aligned). To model the Shift component, we created a *G* matrix with zero entries in the within-start and within-end blocks, and constant value in the start–end cross terms. This model component captures the presence of a uniform, condition-independent shift between start and end postures while preserving internal geometry. Including this Shift component in the regression analysis allowed us to separately quantify the contributions of a shared fixed point geometry (Aligned) and a shift (Shift) to the overall geometry observed in the data.

#### Assessing the alignment of shift dimension

To directly test whether the shift between start and end postures occurred along a consistent neural dimension, we used condition-averaged neural activity from the start and end of each trial (using the same time windows as described above). We randomly selected two of the twenty conditions (without replacement) and computed a shift vector for each by subtracting the end-trial activity from the start-trial activity in the P-dimensional neural space. We then measured the cosine similarity between these two shift vectors. This process was repeated 1,000 times for each brain area and recording session (**Supplementary Fig. 8a**). Across all areas, the shift vectors were consistently aligned, with cosine similarities well above zero—the expected value for randomly oriented vectors in high-dimensional space—indicating a reliable, condition-independent shift direction.

To further confirm that the alignment geometry persists despite the presence of a uniform shift, we tested whether a classifier trained to decode posture location at the start of a trial could generalize to the end of the trial. If the uniform shift disrupted or collapsed the underlying posture geometry, such generalization would not be possible. We performed single-trial classification using logistic regression to predict one of the five posture locations. Neural activity was averaged over a 100 ms window from 150 to 50 ms before target onset for the start of the trial, and over a 200 ms window immediately before trial end for the end of the trial. Classifier performance was evaluated using 10 repetitions of 5-fold cross-validation. As a control, we repeated the same classification procedure with shuffled condition labels to estimate chance-level performance. For each brain region and recording session, we evaluated three settings: (1) train and test on start-of-trial data, (2) train and test on end-of-trial data, and (3) train on start and test on end (**Supplementary Fig. 8b**).

#### Recurrent neural network (RNN) simulations

To examine how the geometry of neural dynamics depends on controller complexity, we used our previously published open-source toolbox, Motornet^87^, which enables training recurrent neural networks (RNNs) to control arm models with arbitrary biomechanical detail. The network received as input the XY coordinates of a target location and a go cue, and was trained to produce control signals that moved an arm model from a random starting position in the workspace to the specified target. We implemented a modular architecture consisting of a PMd-like module of 265 gated recurrent units (GRUs) that received visual feedback from the arm and direct target/go cue input to a 10% subset of units; an M1-like module of 265 GRUs that received input from PMd and minimal (1%) direct target/go cue input; and a spinal cord module of 64 units that integrated input from M1 and proprioceptive signals from the arm, projecting to a fully connected layer that generated muscle or actuator commands. The spinal cord module was included to prevent unrealistically large muscle-like activity in PMd- and M1-like modules. We trained networks with two plant models differing only in biomechanical complexity: a simple point-mass model driven by four linear actuators, and a biomechanically realistic two-joint arm with intersegmental dynamics and six muscles, including shoulder and elbow flexors/extensors and biarticular flexor/extensor pairs. All other aspects of the architecture and training parameters were identical across conditions. We first trained the network on a random reaching task spanning the full workspace with variable delays, then tested it on the five-target layouts used in our monkey experiments. Each trial lasted three seconds, with a variable delay between target onset and the go cue and included no-go (40% in each batch) trials in which the absence of a go cue required the network to maintain its initial position, thereby preventing movement during the preparation period. Networks were trained for 20,000 batches of 128 trials using the Adam optimizer with a learning rate of 1×10^−3^, with the following loss function designed to ensure accurate endpoint placement and smooth movement trajectories.

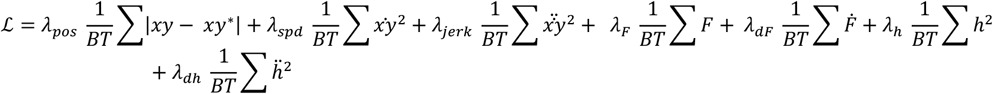

Here, *B* and *T* denote the batch size and number of time steps, respectively. *xy* and *xy*^∗^ are the current and target XY locations. *F* and *T* are the muscle force (or actuator command) and its time derivative, while *h* and 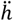 are the GRU hidden state and its second time derivative. The lambda values were set at *λ*_*pos*_*=* 1, *λ*_*spd*_ *=* 1*e* − 3, *λ*_*jerk*_ *=* 1*e* − 4, *λ*_*F*_ *=* 1*e* − 4, *λ*_*dF*_ *=* 1*e* − 4, *λ*_*h*_ *=* 1*e* − 2, *λ*_*dh*_ *=* 1*e* − 1, although precise values were not critical for successful training. Penalties on speed, force, force derivative, and jerk promoted smooth, natural movement^88^. Additional penalties on hidden activity and its temporal derivatives encouraged stable, low-dimensional neural dynamics and discouraged chaotic solutions^89^. Finally, for each plant type we train seven instances of the network and applied the same analysis pipeline used for the monkey data to visualize and quantify the GRU hidden-state trajectories.

## Acknowledgments

This work was supported by a CIHR Project Grant to JD and JAP (PJT-175010) as well as funding from the Azrieli Foundation to JAP (via the Collaboration on Motor Planning, Execution and Resilience). MK was supported by an Ontario Graduate Scholarship. JAM was supported by a Banting Postdoctoral Fellowship, a Vector Institute Postgraduate Affiliation, and a BrainsCAN Postdoctoral Fellowship (Canada First Research Excellence Fund). JAP received a salary award from the Canada Research Chairs Program.

## Author Contribution

Conceptualization: MK, JAP Data collection: MK, JAM, RC

NHP experiment: MK, JAM, RC, JCL, JAP

Analysis and visualization: MK, JAM, JD, JAP RNN

simulations: MK

Supervision: JD, JAP

Writing – original draft: MK

Writing– review and editing: MK, JAM, JD, JAP

## Code and data availability

Neural data analysis and visualization were performed using a custom toolbox (https://github.com/mehrdadkashefi/latent_analysis). All recorded neural data will be available upon publication.

## Supplementary materials

**Supplementary figure 1.**
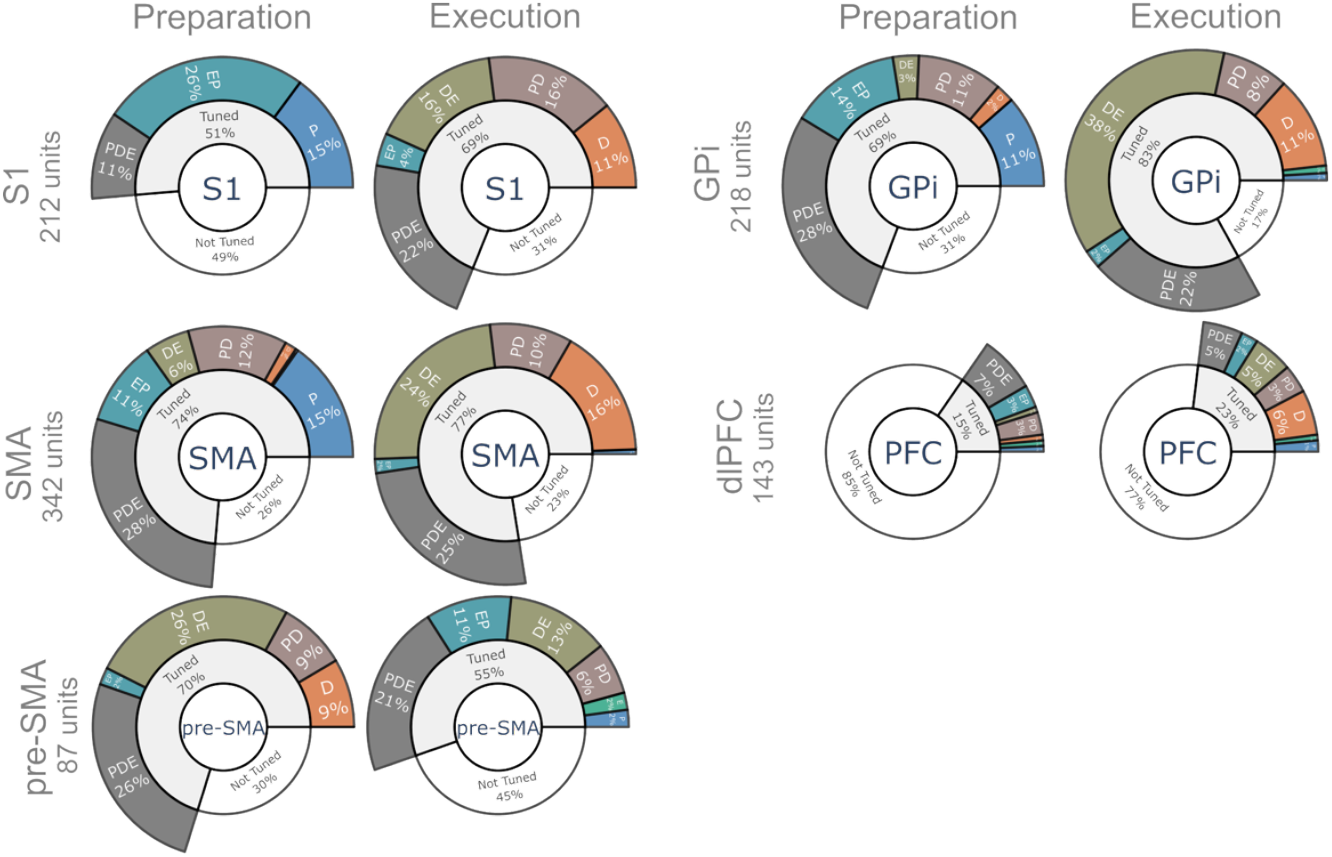
Mixed selectivity in GPi, SMA, preSMA, and dlPFC in Monkey P. Same as Fig. 2B but for all other recorded areas in monkey P.

**Supplementary figure 2.**
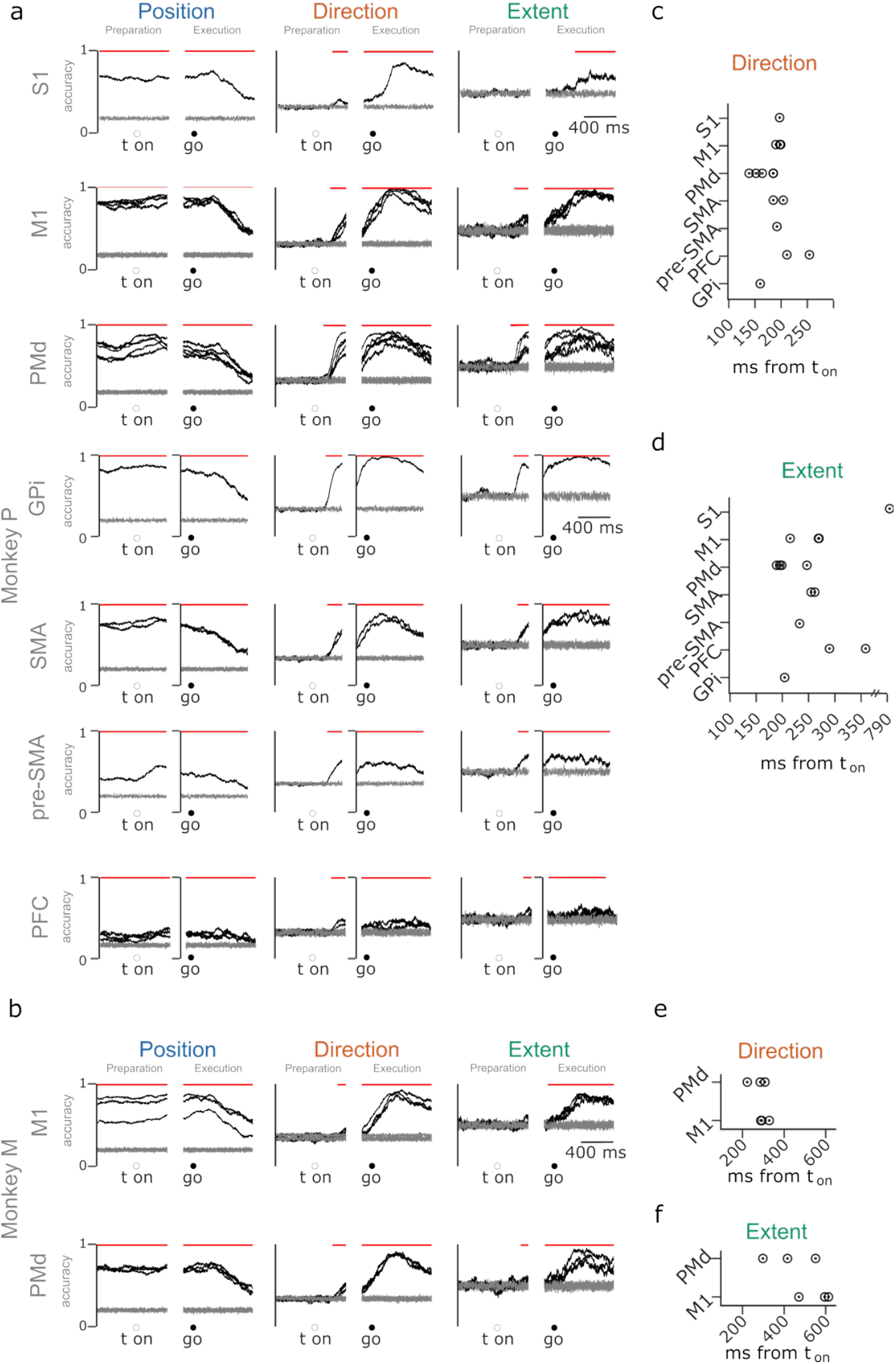
Decoding movement parameters. **a)** Decoding of movement position, direction, and extent throughout the trial from the firing rates of all recorded areas in Monkey P. Neural data were aligned to both target onset (t on) and the go cue (go). Black traces indicate decoding accuracy; gray traces represent chance level. Each trace corresponds to one recording session. Red lines mark time points where accuracy was significantly above chance. **b)** Same as (a) but for Monkey M. **c)** First time point relative to target onset when movement direction was significantly decodable across all areas in Monkey P. **d)** Same as (c) but for movement extent in Monkey P. **e)** Same as (c) but for Monkey M. **f)** Same as (d) but for movement extent in Monkey M.

**Supplementary figure 3.**
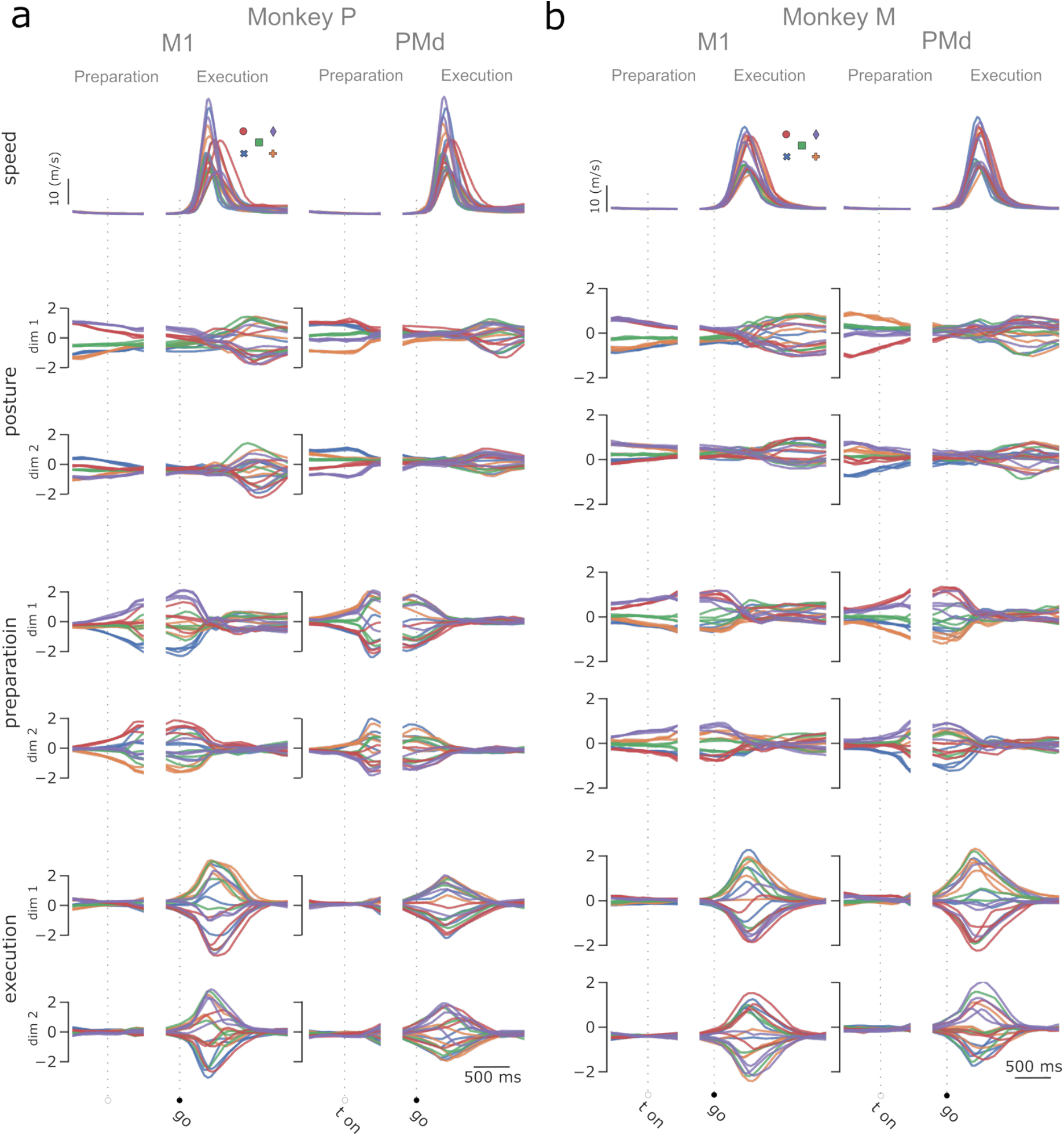
Projection of neural data on isolated posture, preparation and execution dimension. **a)** Speed profiles and subspace projections for Monkey P. Top row: Average speed profiles. Rows 2–7: Projections of M1 and PMd neural activity onto the top two dimensions of the posture (rows 2–3), preparation (rows 4–5), and execution (rows 6–7) subspaces. Neural activity is aligned to target onset for the preparation phase and to the go cue for the execution phase. Traces are color-coded according to the location of the home target. **b)** Same as (a) for Monkey M

**Supplementary figure 4.**
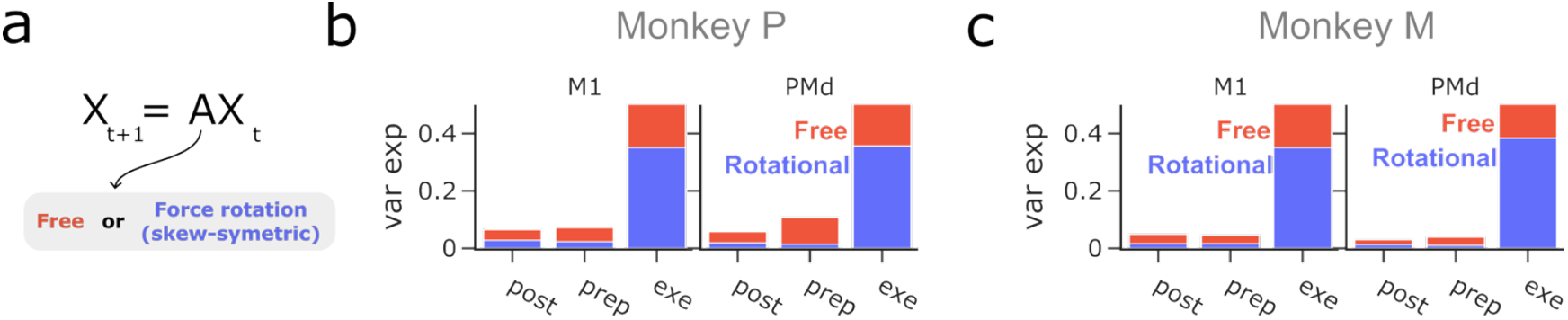
Strength of rotational dynamics in isolated subspaces. **a)**Execution-phase neural activity (aligned to the go cue) was projected onto the posture, preparation, and execution subspaces, followed by the fitting of a linear dynamical model. The model utilized either an unconstrained transition matrix (*“*free*”*) or a transition matrix constrained to be skew-symmetric, which forces purely rotational dynamics. **b)** Variance explained by the free (red) and forced-rotational (blue) models for each subspace (posture, preparation, and execution) in M1 and PMd for Monkey P. **c)** Same as (b) but for Monkey M.

**Supplementary figure 5.**
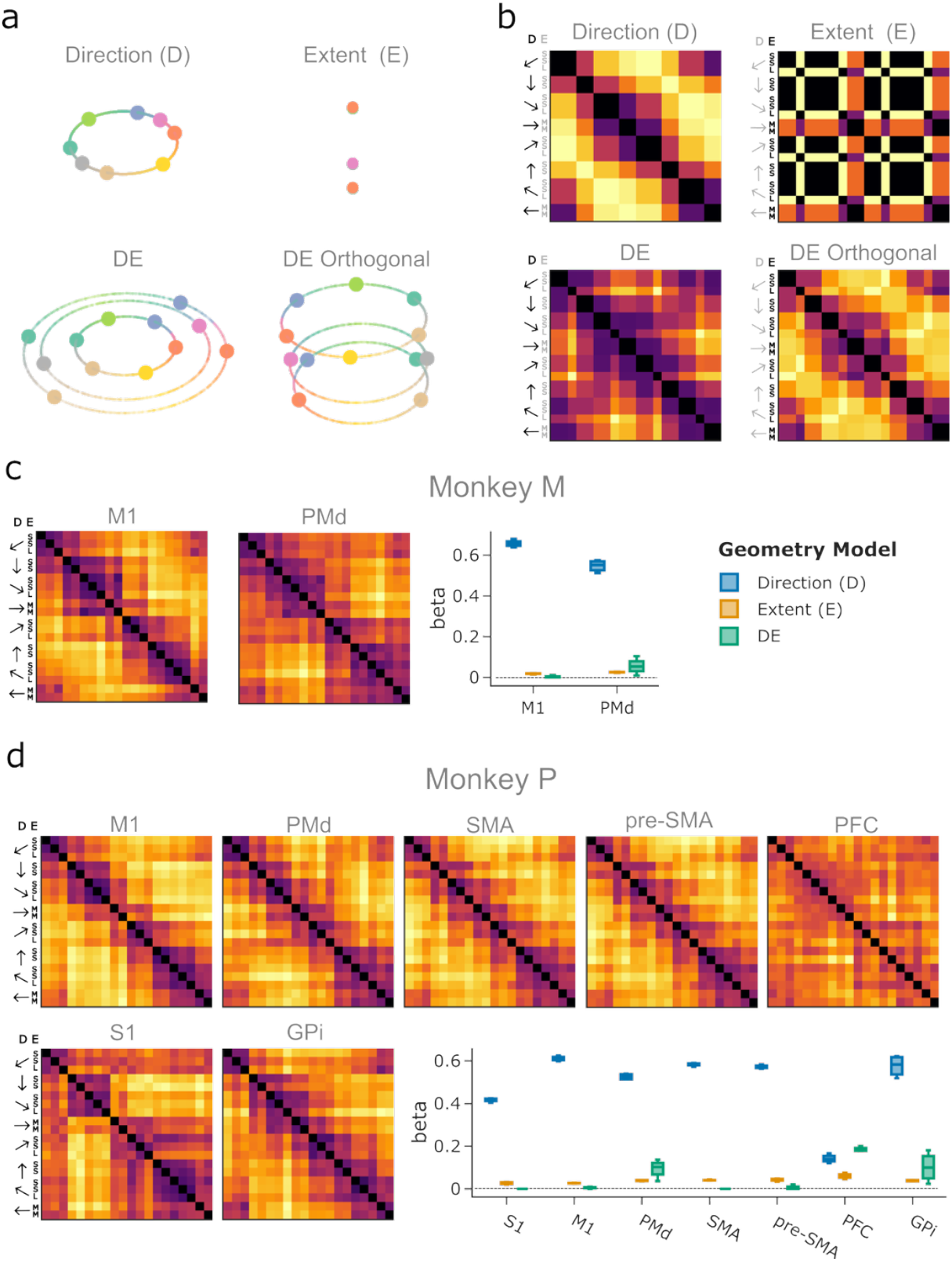
Geometry of neural state space at peak speed. **a)** Schematic of four hypothetical geometrical arrangements of movement direction and extent in neural state space at peak speed. In the Direction-only (D), Direction-Extent (DE), and Direction-Extent Orthogonal (DE-Orthogonal) models, movement direction is represented along the circumference of a ring. In the Extent-only (E) model, extent is encoded along a linear axis. In the DE model, extent is represented by the radius of concentric rings; in the DE-Orthogonal model, extent is encoded along an axis orthogonal to the plane containing the directional rings. **b)** Representational dissimilarity matrices (RDMs) corresponding to each of the four models in (a). **c)** RDM derived from neural data at peak movement speed in M1 and PMd of Monkey M, along with non-negative regression beta weights indicating each model*’*s contribution to the observed neural geometry. **d)** Same as (c), but for all recorded brain regions in Monkey P.

**Supplementary figure 6.**
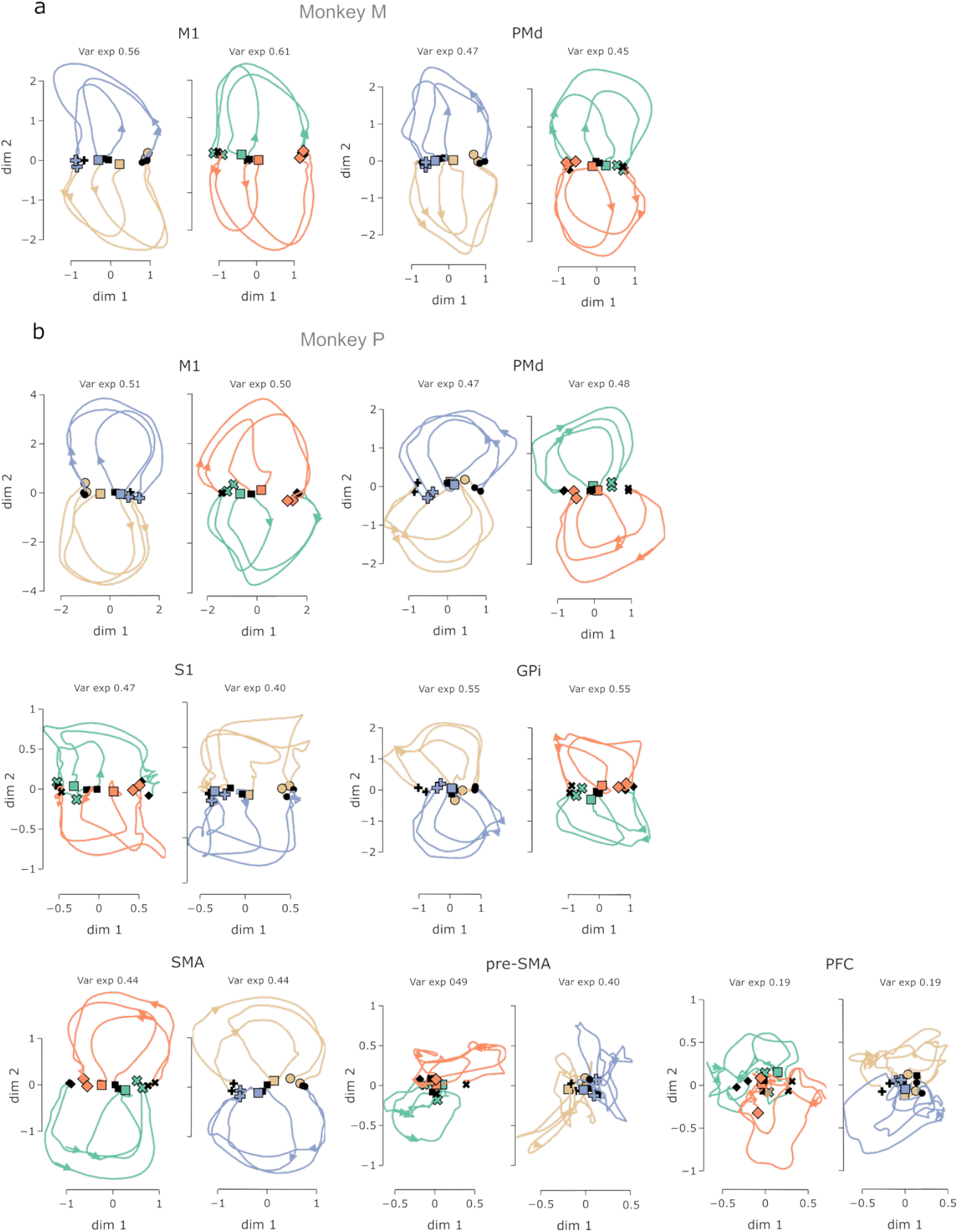
Rotational dynamics connect posture-specific fixed points during reaching. **a)** Same as Fig. 3A–B, but for M1 and PMd in Monkey M. **b)** Same as A, but for all recorded areas in Monkey P.

**Supplementary figure 7.**
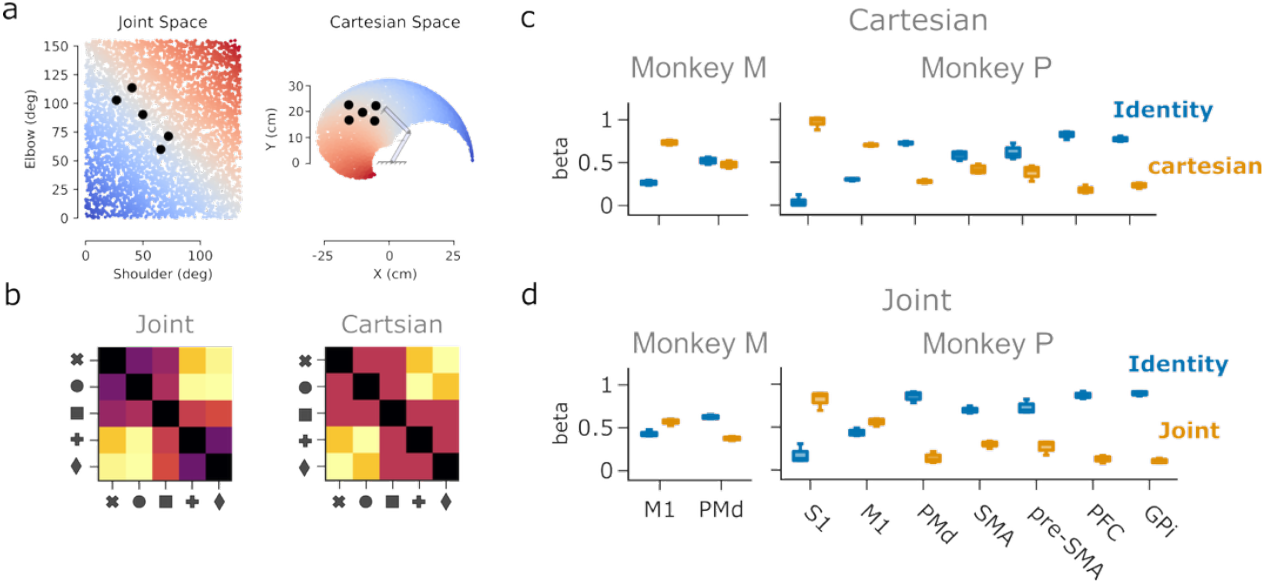
Replicating the analysis in figure 4 with joint geometry. **a)** Joint and Cartesian spaces for a uniformly sampled two-dimensional workspace of a two-joint arm. Sample locations are colored from fully flexed (blue) to fully extended (red) arm configurations. Black dots indicate the approximate positions of the five target locations used in the NHP experiment. **b)** Representational dissimilarity matrices (RDMs) for the distances between the five selected target locations in joint space (left) and Cartesian space (right). **c)** Same as Fig. (4c–d), showing the contributions of the Identity and Cartesian models. **d)** Same as (c), but with the Joint model in place of the Cartesian model.

**Supplementary figure 8.**
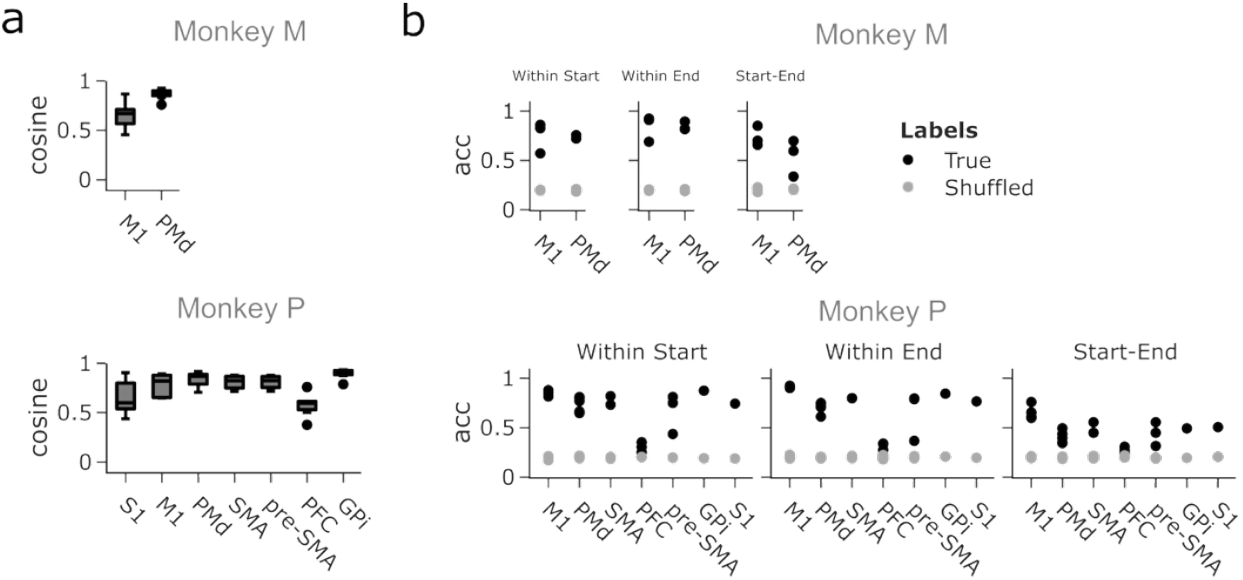
Alternative methods characterizing the uniform shift. **a)** Uniformity of shift vectors: mean cosine similarity values for shift vectors computed from randomly sampled pairs of conditions (see Methods). **b)** Alignment between start and end epochs: classification accuracy for decoders trained to discriminate posture locations, either trained and tested within the start epoch (*Within Start*), trained and tested within the end epoch (*Within End*), or trained on the start epoch and tested on the end epoch (*Start–End*). Gray dots indicate accuracy from classifiers with shuffled labels (chance level).

**Supplementary figure 9.**
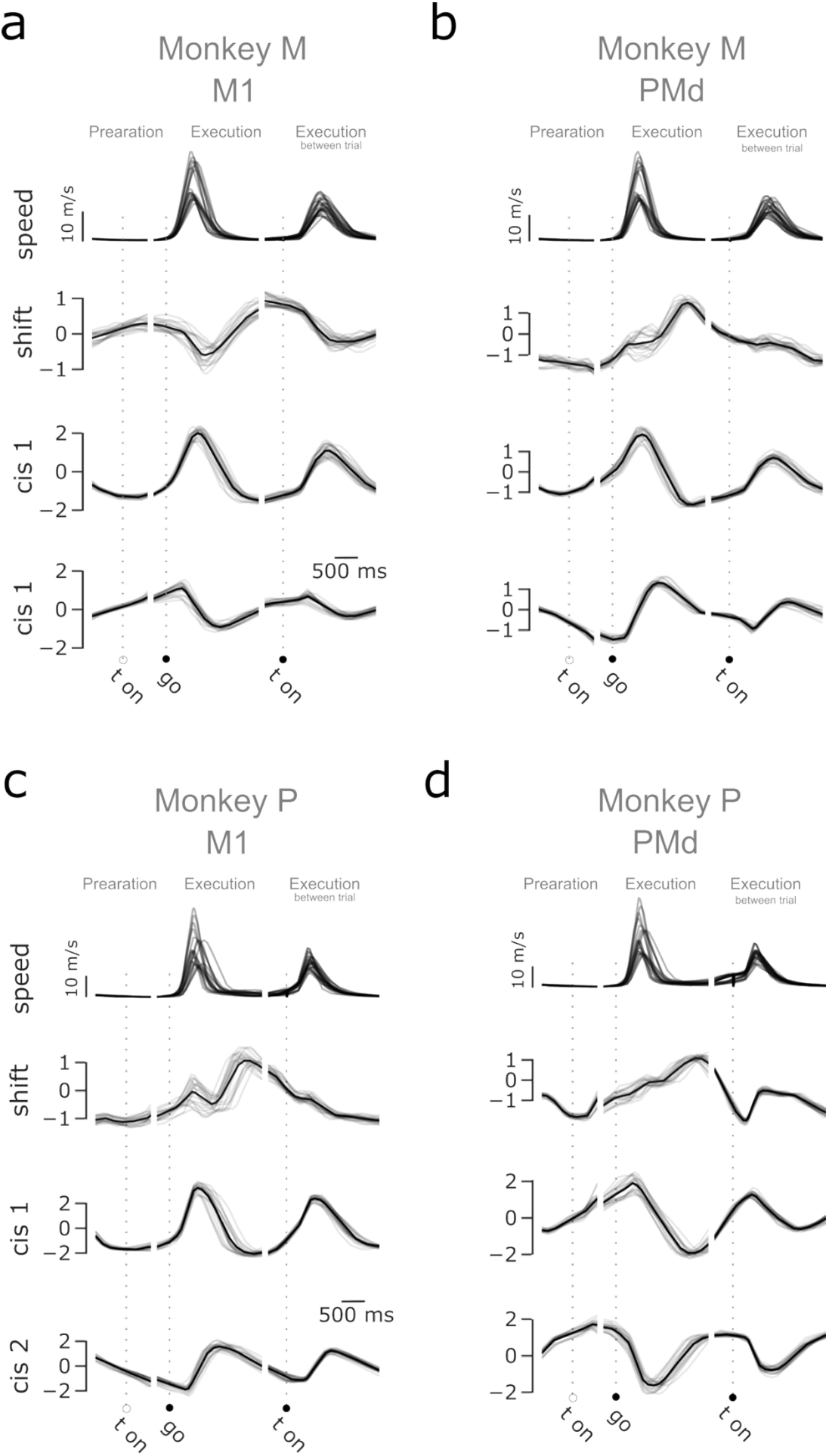
Comparison between condition independent signal (CIS), and uniform shift dimension. **a)** Average speed profiles and neural projections for Monkey M (M1). Top row: Average speed profiles. Second row: Projection of M1 activity onto the uniform shift dimension during preparation, execution, and inter-trial movements. Third and fourth rows: Projections onto the top two condition-independent signals (CIS 1 and CIS 2). Individual conditions are shown in gray, with the cross-condition mean indicated by the bold dark line. Data are shown across preparation, execution, and the between-trial interval. **b)** Same as (a) but for Monkey M*’*s PMd. **c)** Same as (a) for Monkey P*’*s M1. **d)** Same as (b) for Monkey P*’*s PMd.

**Supplementary table 1.** List of recording sessions.

| Monkey | Session | Successful trials | Recording area(s) | Recorded neurons (KS good) |
| --- | --- | --- | --- | --- |
| M | May 09 2024 | 631 | M1 | 339 |
|  | May 10 2024 | 857 | M1 | 267 |
|  | May 13 2024 | 634 | PMd | 273 |
|  | May 14 2024 | 415 | PMd | 303 |
|  | May 15 2024 | 797 | PMd | 257 |
|  | May 16 2024 | 907 | M1 | 347 |
| P | May 13 2024 | 641 | M1 | 404 |
|  | May 14 2024 | 605 | M1-PMd | 252-310 |
|  | May 15 2024 | 777 | M1-PMd | 237-178 |
|  | May 16 2024 | 634 | GPI | 549 |
|  | May 17 2024 | 641 | PMd | 163 |
|  | May 21 2024 | 475 | M1-dLPFC | 333-151 |
|  | May 22 2024 | 775 | dLPFC | 132 |
|  | May 23 2024 | 789 | SMA-preSMA | 303-125 |
|  | May 24 2024 | 635 | SMA-dLPFC | 290-152 |
|  | June 20 2025 | 964 | PMd | 132 |
|  | June 21 2025 | 775 | S1-PMd | 240-245 |

